# Massively parallel combination screen reveals small molecule sensitization of antibiotic-resistant Gram-negative ESKAPE pathogens

**DOI:** 10.1101/2024.03.26.586803

**Authors:** Megan W. Tse, Meilin Zhu, Benjamin Peters, Efrat Hamami, Julie Chen, Kathleen P. Davis, Samuel Nitz, Juliane Weller, Thulasi Warrier, Diana K. Hunt, Yoelkys Morales, Tomohiko Kawate, Jeffrey L. Gaulin, Jon H. Come, Juan Hernandez-Bird, Wenwen Huo, Isabelle Neisewander, Laura L. Kiessling, Deborah T. Hung, Joan Mecsas, Bree B. Aldridge, Ralph R. Isberg, Paul C. Blainey

**Author notes:** These authors contributed equally (Megan W. Tse, Meilin Zhu). These authors contributed equally (Benjamin Peters, Efrat Hamami). These authors are co-corresponding and contributed equally (Ralph R. Isberg, Paul C. Blainey). **Corresponding authors**: Paul C. Blainey **Email:**, Ralph R. Isberg **Email:**. **Author Contributions:** M.W.T., M.Z., B.P., E.H., J.C., K.P.D., J.W., T.W., L.L.K., D.T.H., J.M., B.B.A., R.R.I., and P.C.B. designed research. M.W.T., M.Z., B.P., J.C., K.P.D., and J.W. performed research. E.H., S.N., T.W., D.H., Y.M., T.K., J.G., J.H.C., J.H., W.H., and I.N. contributed new reagents or analytic tools. M.W.T., M.Z., B.P., E.H., K.P.D., J.C., S.N, and J.W. analyzed data. M.W.T., M.Z., R.R.I., and P.C.B. wrote the paper. **Competing Interests Statement:** P.C.B. is a consultant to or holds equity in 10X Genomics, General Automation Lab Technologies/Isolation Bio, Celsius Therapeutics, Next Gen Diagnostics, Cache DNA, Concerto Biosciences, Stately, Ramona Optics, Bifrost Biosystems, and Amber Bio. His laboratory has received research funding from Calico Life Sciences, Merck, and Genentech for unrelated work. The Broad Institute and MIT may seek to commercialize aspects of this work, and related applications for intellectual property have been filed. **Classification:** Biological Sciences, Microbiology.

## Abstract

Antibiotic resistance, especially in multidrug-resistant ESKAPE pathogens, remains a worldwide problem. Combination antimicrobial therapies may be an important strategy to overcome resistance and broaden the spectrum of existing antibiotics. However, this strategy is limited by the ability to efficiently screen large combinatorial chemical spaces. Here, we deployed a high-throughput combinatorial screening platform, DropArray, to evaluate the interactions of over 30,000 compounds with up to 22 antibiotics and 6 strains of Gram-negative ESKAPE pathogens, totaling to over 1.3 million unique strain-antibiotic-compound combinations. In this dataset, compounds more frequently exhibited synergy with known antibiotics than single-agent activity. We identified a compound, P2-56, and developed a more potent analog, P2-56-3, which potentiated rifampin (RIF) activity against *Acinetobacter baumannii* and *Klebsiella pneumoniae*. Using phenotypic assays, we showed P2-56-3 disrupts the outer membrane of *A. baumannii*. To identify pathways involved in the mechanism of synergy between P2-56-3 and RIF, we performed genetic screens in *A. baumannii*. CRISPRi-induced partial depletion of lipooligosaccharide transport genes (*lptA*-*D*, *lptFG*) resulted in hypersensitivity to P2-56-3/RIF treatment, demonstrating the genetic dependency of P2-56-3 activity and RIF sensitization on *lpt* genes in *A. baumannii.* Consistent with outer membrane homeostasis being an important determinant of P2-56-3/RIF tolerance, knockout of maintenance of lipid asymmetry complex genes and overexpression of certain resistance-nodulation-division efflux pumps – a phenotype associated with multidrug-resistance – resulted in hypersensitivity to P2-56-3. These findings demonstrate the immense scale of phenotypic antibiotic combination screens using DropArray and the potential for such approaches to discover new small molecule synergies against multidrug-resistant ESKAPE strains.

**Significance Statement:** There is an unmet need for new antibiotic therapies effective against the multidrug-resistant, Gram-negative ESKAPE pathogens. Combination therapies have the potential to overcome resistance and broaden the spectrum of existing antibiotics. In this study, we use DropArray, a massively parallel combinatorial screening tool, to assay more than 1.3 million combinations of small molecules against the Gram-negative ESKAPE pathogens, *Acinetobacter baumannii*, *Klebsiella pneumoniae*, and *Pseudomonas aeruginosa*. We discovered a synthetic small molecule potentiator, P2-56, of the antibiotic rifampin effective in *A. baumannii* and *K. pneumoniae*. We generated P2-56-3, a more potent derivative of P2-56, and found that it likely potentiates rifampin by compromising the outer membrane integrity. Our study demonstrates a high-throughput strategy for identifying antibiotic potentiators against multidrug-resistant bacteria.

## Introduction

Antibiotic resistance is an ongoing and increasing worldwide problem. In 2019, antibiotic-resistant bacterial infections led to 1.27 million deaths globally (1). By 2050, experts estimate such infections will cause up to 10 million deaths per year globally (2). With the growing concerns of antibiotic resistance, the World Health Organization designated a set of priority pathogens for the discovery and development of new antibacterials. These pathogens, collectively known as the ESKAPE pathogens, encompass *Enterococcus faecium*, *Staphylococcus aureus*, *Klebsiella pneumoniae*, *Acinetobacter baumannii*, *Pseudomonas aeruginosa*, and *Enterobacter* spp. (3). Despite the impact of these antibiotic-resistant organisms, the rate of FDA-approvals for new antibacterials has diminished markedly over the past 50 years (4). Of the antibiotic drug approvals in the last 10 years, none exhibited new mechanisms of action, rendering these drugs potentially liable to existing widespread resistance mechanisms (5). Thus, there exists a critical need for new strategies that identify clinically useful antimicrobial activities against the Gram-negative ESKAPE pathogens: *A. baumannii*, *K. pneumoniae*, and *P. aeruginosa* (3, 6).

The most successful antibiotics with reduced rates of clinical resistance have been observed to possess activity against more than one target, referred to as multi-targeting (7). However, the discovery and development of new single-agents with multi-targeting effects has been challenging. Thus, combinations of small molecules may serve as an alternate strategy to address multiple essential targets or generate synthetic lethal effects (8, 9). Combinations of small molecules have already emerged as an important strategy in addressing the antibiotic resistance crisis, primarily by targeting resistance factors known to compromise the activity of a single antibiotic (9). For example, beta-lactamase inhibitors, currently used in the clinic in combination with beta-lactam antibiotics, emerged from directed efforts to restore beta-lactam efficacy (10, 11). Other commonly prescribed antibiotic combinations, such as trimethoprim-sulfamethoxazole, were discovered independently and combined post-hoc due to their increased efficacy when used in combination (12). Thus, combinations can open opportunities to overcome an unrecognized resistance mechanism via synergistic activity and/or increase an existing antibiotic’s potency (8, 9, 13). We propose high-throughput phenotypic combinatorial screening to discover such antibiotic combinations. However, large-scale discovery of new drug interactions using conventional screening approaches is challenging, as even small increases in the number of drugs being tested can demand an intractable number of assays. Thus, we urgently need new methods to efficiently discover efficacious antibiotic combinations.

Precise liquid handling operations and high compound consumption limit conventional combinatorial drug screens. Consequently, conventional plate-based antibiotic combination screens have prioritized testing smaller panels of antibiotics (up to 5) screened in combination with large compound libraries against a single pathogen strain (14–16). The droplet-in-microarray technology, referred to as DropArray, demonstrated its ability to efficiently screen combinatorial chemical spaces across both antibiotic and compound axes, identifying synergistic drug combinations across 4,000 repurposing drugs and 10 antibiotics against *E. coli* (17). Through nanoliter-scale miniaturization and random self-assembly of droplet combinations on a microwell array chip, DropArray circumvents the limitations of standard plate-based methods (17). Thus, DropArray-based combinatorial screening presents an opportunity to discover potentiators that enhance antibiotic efficacy for clinically-relevant Gram-negative ESKAPE pathogens.

In this work, we scaled up DropArray using a panel of 6 Gram-negative ESKAPE pathogen strains, up to 22 antibiotics, and over 30,000 chemical compounds (Fig. 1A,B; Table S1) to orient large-scale combinatorial screening efforts towards clinically-relevant pathogens. While all 6 strains show significant resistances, we selected at least one highly resistant clinical isolate within each species (Fig. 1B; Table S1) with the expectation that this would reduce the hit rate and apparent potency of hit compounds and enrich the hit set for combinations able to suppress the growth of resistant ESKAPE strains and compound scaffolds able to evade or attack important extant mechanisms of antibiotic resistance. Through DropArray-based combinatorial screening, we discovered P2-56, a small molecule that synergized with an assortment of Gram-positive-acting antibiotics, including rifampin (RIF), against *A. baumannii* and *K. pneumoniae*. We demonstrated using plate-based phenotypic assays that a more potent analog, P2-56-3, increased the outer membrane permeability in *A. baumannii*. Using CRISPRi-induced hypomorphic gene depletions in *A. baumannii*, we identified the knockdown of the lipooligosaccharide (LOS) transport pathway conferred hypersensitivity to P2-56-3/RIF and P2-56-3 treatments. Additionally, we found that outer membrane-compromising defects, such as the maintenance of lipid asymmetry (MLA) protein complex and resistance-nodulation-division (RND) pump overexpression, caused hypersensitivity to P2-56-3/RIF treatment in *A. baumannii*. Together, this work demonstrates the utility of DropArray in phenotypic identification of promising antibiotic combinations against the Gram-negative ESKAPE pathogens and genetic screens to efficiently probe the genetic dependencies of synergistic drug interactions.

**Figure 1.**
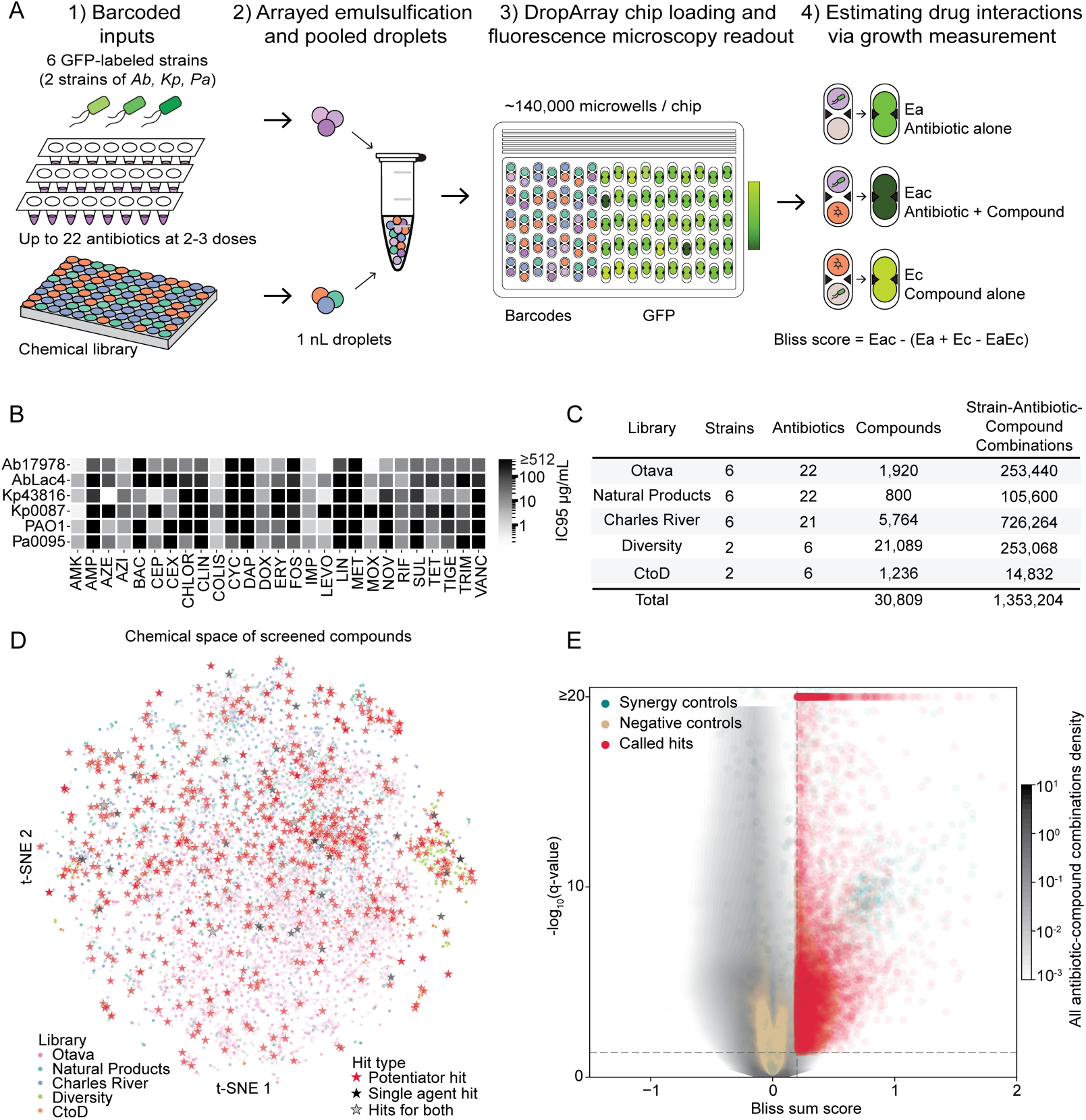
Overview of the antibiotic combinations screen using the DropArray platform. (A) The workflow for antibiotic combinations screening in DropArray. The workflow for antibiotic combinations screening in DropArray. Each microwell carried two droplets where a droplet contained either GFP-expressing strains with antibiotics at 2–3 concentrations or compounds from chemical libraries. We 1) encoded assay inputs with fluorescent barcodes, 2) emulsified them into nanoliter droplets and pooled for 3) loading and random self-assembly of pairwise combinations on the discoveryChip. We imaged microwells prior and after droplet merging, and 4) measured GFP fluorescence as a proxy for bacterial biomass. (B) The panel of Gram-negative pathogen strains and antibiotics screened. In the heatmap, we indicated the approximate IC90 (µg/mL) determined in plates for each antibiotic and strain (Supplemental Methods). (C) Number of strains, antibiotics, compounds and strain-antibiotic-compound combinations screened per library. (D) Chemical space depicted using a t-SNE of the Morgan fingerprints for each compound screened for antibiotic potentiating activity. We included one-third of all compounds that passed screening quality metrics for representation (Supplemental Methods). Solid red stars represent potentiator hits (compounds with a Bliss sum score ≥0.3 and FDR q-values ≤0.05 with at least one strain-antibiotic pair), solid black stars represent single-agent hits (compounds exhibiting ≥30% growth inhibitory activity with FDR q-values ≤0.05 in at least one strain), and black-outlined gray stars represent compounds scoring as both a potentiator and a single-agent hit. (E) Bliss sum scores and significance values for all strain-antibiotic-compound combinations screened (black), called combination hits (red), selected antibiotic-antibiotic positive controls (green), and antibiotic-media negative controls (beige) represented as a volcano plot (hit-calling thresholds: Bliss sum score ≥0.3 and FDR q-values ≤0.05). P-value and FDR calculations: Materials and Methods.

## Results

### Antibiotic combination discovery across the Gram-negative ESKAPE pathogens

We leveraged DropArray to identify new antibiotic combinations that were synergistic against 6 Gram-negative ESKAPE pathogen strains (Fig. 1A) with diverse antibiotic susceptibility profiles: *A. baumannii* ATCC 17978, *A. baumannii* LAC-4, *K. pneumoniae* ATCC 43816, *K. pneumoniae* AR0087, *P. aeruginosa* PAO1, and *P. aeruginosa* AR0095 (Fig. 1B; Table S1; Materials and Methods). Within each species, we selected a single clinical isolate strain that broadly exhibited high resistance towards the antibiotic screening panel (Fig. 1B; Table S1). To measure growth inhibitory effects in the DropArray platform (Fig. S1A,B), we engineered each of these strains to express GFP and used fluorescence as a proxy for bacterial biomass (Materials and Methods). Our antibiotic panels included up to 22 antibiotics with diverse mechanisms, each tested at 2–3 concentrations (Fig. S2A). In combination with these antibiotics, we assayed: investigational synthetic compounds from the Otava, Charles River, and Diversity-Oriented Synthesis (Diversity) libraries (18, 19); semi-synthetic compounds from the Complex-to-Diversity (CtoD) collection (20, 21); and natural product compounds from the Microsource Discovery Systems Natural Products library. To identify the antibiotic and compound inputs of each assay, we used a previously described fluorescent dye-based barcoding strategy for DropArray assays (Fig. 1A; Materials and Methods) (17, 22). To measure the data quality of each chip, we calculated Z-prime scores using GFP fluorescence values from the bacteria only and media only combinations as our positive and negative assay controls, respectively. Finally, we designed our screens to have a median of 10 or more replicates per combination, targeting a Z-prime cutoff of 0.2 for controls (Fig. S2B; Materials and Methods; Supplemental Methods).

We tested over 30,000 unique compounds in combination with a panel of up to 22 antibiotics against 2 or 6 strains of the priority Gram-negative ESKAPE pathogens (Fig. 1C; Table S2). In total, we screened over 81 million microwell assays and over 1.3 million unique strain-antibiotic-compound combinations, with a median replication level of 15 (Fig. 1C; Table S2). After excluding chips failing to meet a Z-prime cutoff of 0.2, our dataset included 85% of the total chips screened (Table S2; Materials and Methods) and represented a diverse chemical space (Fig. 1D). Because the DropArray platform uses self-assembly to formulate all possible combinations of inputs, our screen produced measurements for the independent effect of each antibiotic and each compound in addition to the combined effect of antibiotic-compound combination for all strains. We estimated drug interactions using the Bliss score, determined by subtracting the independent effects of the antibiotic and of the compound from the combination growth inhibition effect (Fig. 1A; Materials and Methods) (23). Thus, DropArray can support the discovery of both synergistic drug interactions between antibiotics and compounds and of single-agent activity of each compound against the Gram-negative ESKAPE pathogens.

### Screened compounds more commonly have synergistic drug interactions with known antibiotics than single-agent activity

To summarize composite small molecule combination interaction scores, we calculated a single, cumulative interaction statistic by summing the individual Bliss scores across the tested antibiotic concentrations for each antibiotic-compound combination, referred to as the Bliss sum score. We selected antibiotic pairs with previously documented and/or experimentally validated synergistic activity as positive synergy controls (Fig. S2C; Table S2) (11, 12) and antibiotic-media pairs that have no expected drug interactions as negative synergy controls. We used these positive and negative synergy controls to determine hit-calling thresholds such that the majority of the positive controls met the criteria used to define hits and over 99.66% of the negative controls did not meet the criteria used to define hits (Bliss sum score ≥0.3 and FDR q-value ≤0.05; Fig. 1E; Dataset S1, S2). We note that these cutoffs resulted in a lower true positive rate of the positive synergy controls for the CtoD library screen, due to the selected antibiotic concentrations (Fig. S2C).

The selected thresholds resulted in 3,784 antibiotic-compound combination synergy hits spanning 2,388 unique library compounds. For individual compound activity with each strain, we identified 244 total single-agent hits spanning 129 unique library compounds (growth inhibition ≥30% and FDR q-value ≤0.05; Dataset S3). In the screened chemical space, we observed the combination hit rate (8.57%, 2388/27,854) for library compounds exceeded the single-agent hit rate by more than order of magnitude (0.46%, 129/27,854; Fig. 1D; Supplemental Text) indicating the increased potential for discovering new antimicrobial strategies through combinatorial screening. Additionally, the DropArray platform provides a comprehensive activity profile for each screened compound including its independent effect and interaction effects with all strains and antibiotics, enabling interpretation to support hit prioritization.

### Synergistic interaction hits validate in checkerboard assays

To confirm the synergistic effects observed in the primary screen, we evaluated select antibiotic-compound combinations using checkerboard interaction assays in DropArray (unless otherwise noted) with greater dose range and resolution. We resupplied and retested 158 compound potentiator hits after assessing their Bliss sum scores from the primary screen, compound availability, and prioritizing compounds that exhibited synergistic activity with multiple antibiotics and/or strains. We called combinations as synergistic in checkerboard assays if a Bliss sum score equalled or exceeded 0.3 across retested antibiotic concentrations with the same compound concentration used in the initial screen. By this criterion, 52% (82/158) of the resupplied compound potentiator hits showed synergistic drug interactions again in at least one antibiotic-strain condition (Fig. S3A). Additionally, 93% (25/27) of the resupplied compounds tested for their single-agent activity demonstrated validated growth inhibitory activity (≥30% growth inhibition at 50 µM against at least one strain; Fig. S4). Unsurprisingly, compounds that are known antibiotics, such as salinomycin (24), hygromycin B (25), and kasugamycin hydrochloride (26), exhibited strong single-agent growth inhibitory activity (Dataset S3). The recovery of compounds with known activity provided confidence in the sensitivity of the DropArray platform and screen.

From these checkerboard assays, we highlight select compounds that showed strong synergistic interactions with at least one strain-antibiotic condition, including a range of combination activities that validated against highly antibiotic-resistant strains (Table S3). The following combinations exhibited a minimum fractional inhibitory concentration ≤0.5 – a different and stringent measure of synergy (Supplemental Methods). BRD0433, from the Diversity collection, demonstrated synergy with RIF against *K. pneumoniae* ATCC 43816 (Fig. S3B) and AR0087 (Table S3) in checkerboard assays. IIIA:3:G, a derivative of abietic acid from the CtoD set, showed synergy with RIF against *K. pneumoniae* ATCC 43816 (Fig. S3C) and AR0087 (Table S3). Interestingly, BRD1479, which is structurally similar to sulfamethoxazole, demonstrated its synergistic interactions with trimethoprim against *P. aeruginosa* PAO1 (Fig. S3D) and AR0095 (Table S3). The structural similarity between BRD1479 and sulfamethoxazole suggested a similar mechanism of synergy as that described for the clinically used combination treatment sulfamethoxazole and trimethoprim, which blocks two different proteins involved in folate synthesis (12). BRD4550, another sulfonamide-like structure, also showed synergy with β-lactams, cefepime, ampicillin, and imipenem in the primary screen and demonstrated synergy with cefepime in a checkerboard assay against *A. baumannii* ATCC 17978 (Fig. S3E) and *P. aeruginosa* AR0095 (Table S3). From the natural products library, esculin monohydrate demonstrated synergy with azithromycin against *K. pneumoniae* AR0087 in checkerboard assays (Fig. S3F). Thus, DropArray’s ability to reproduce compounds with expected synergistic interactions, in addition to growth inhibitory effects, further established confidence in the platform’s ability to sensitively measure growth inhibition and drug interactions.

Notably, we found that P2-56, an indole piperidine from the Otava library, potentiated the effect of multiple antibiotics within *A. baumannii* and *K. pneumoniae* strains and at least one antibiotic across all the Gram-negative ESKAPE species (Fig. 2A,B). This specific compound to our knowledge has not been previously reported to synergize with any antibiotics used against the Gram-negative ESKAPE pathogens. P2-56 exhibited synergy primarily with Gram-positive-acting antibiotics, such as rifampin (RIF), novobiocin, and clindamycin against all *A. baumannii* and *K. pneumoniae* strains included in the primary screen (Fig. 2B). To ensure activity was derived from the pure compound annotated in the commercial library, we resynthesized a racemic mixture of P2-56 and evaluated its activity with RIF against *A. baumannii*, *K. pneumoniae*, and *P. aeruginosa* strains in microtiter plates. This resynthesized compound maintained synergistic activity with RIF in *A. baumannii* and *K. pneumoniae* (Fig. 2C-F). Additionally, the P2-56/RIF combination treatment demonstrated its highest effectiveness in *A. baumannii*, and particularly in the more drug-resistant LAC-4 strain (Fig. 2C,D). Thus, we proceeded to focus on P2-56/RIF synergy in *A. baumannii*.

**Figure 2.**
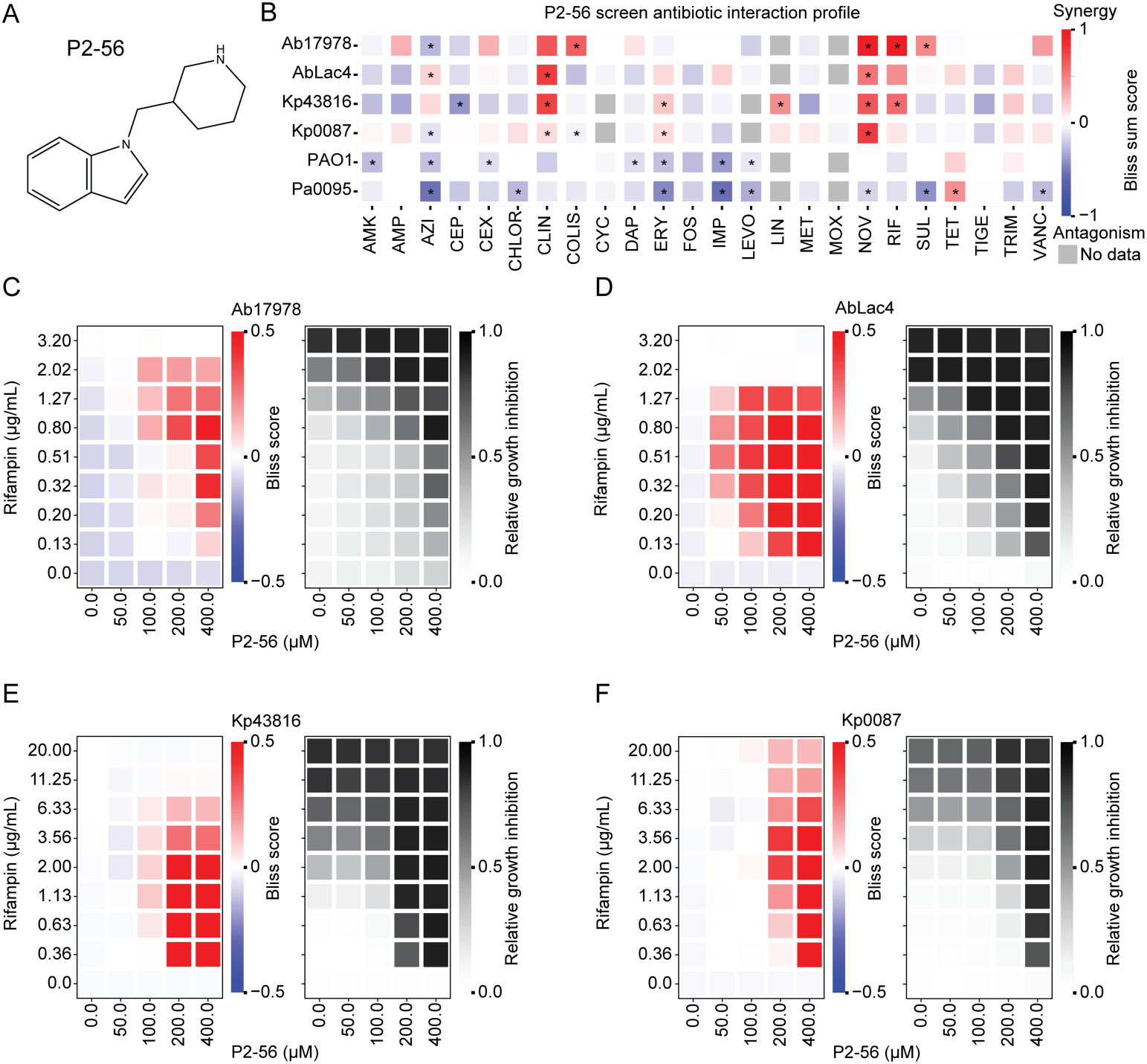
P2-56 synergizes with RIF in *A. baumannii* and *K. pneumoniae.* (A) Chemical structure of P2-56. (B) Antibiotic interaction profile of P2-56 from the primary DropArray screening data where we estimated interactions using Bliss sum scores (synergy in red, antagonism in blue, and not tested in gray). *: FDR q-value ≤0.05. Bliss scores and growth inhibition values from plate-based checkerboards of resynthesized P2-56 and RIF in (C) *A. baumannii* ATCC 17978, (D) *A. baumannii* LAC-4, (E) *K. pneumoniae* ATCC 43816, and (F) *K. pneumoniae* AR0087 (values represent the mean of 2 technical replicates and 2 biological replicates). P-value and FDR calculations: Materials and Methods.

### P2-56-3 synergizes with RIF across multiple Gram-negative strains

To explore the synergistic potency of the P2-56 scaffold with RIF against *A. baumannii*, we synthesized and tested a series of structural analogs of P2-56 in combination with RIF across a diverse panel of *A. baumannii* strains. These strains included the screening strain ATCC 17978 and the following clinical isolates: AB5075 (bone isolate) (27), EGA355 (sputum isolate), EGA366 (urine isolate), and EGA368 (sputum isolate). We identified P2-56-3, a structural analog containing an isomeric piperidine, that was more potently synergistic with RIF across all 5 strains of *A. baumannii* relative to the parent compound, P2-56 (Fig. 3A,B). To evaluate the interaction of P2-56-3 with other Gram-positive-acting antibiotics from the primary screen, we tested P2-56-3 in combination with novobiocin, linezolid, erythromycin, and vancomycin against *A. baumannii* using checkerboard assays in microtiter plates. We observed synergistic effects with novobiocin, linezolid, and erythromycin but not vancomycin (Fig. 3C; S5A-C).

**Figure 3.**
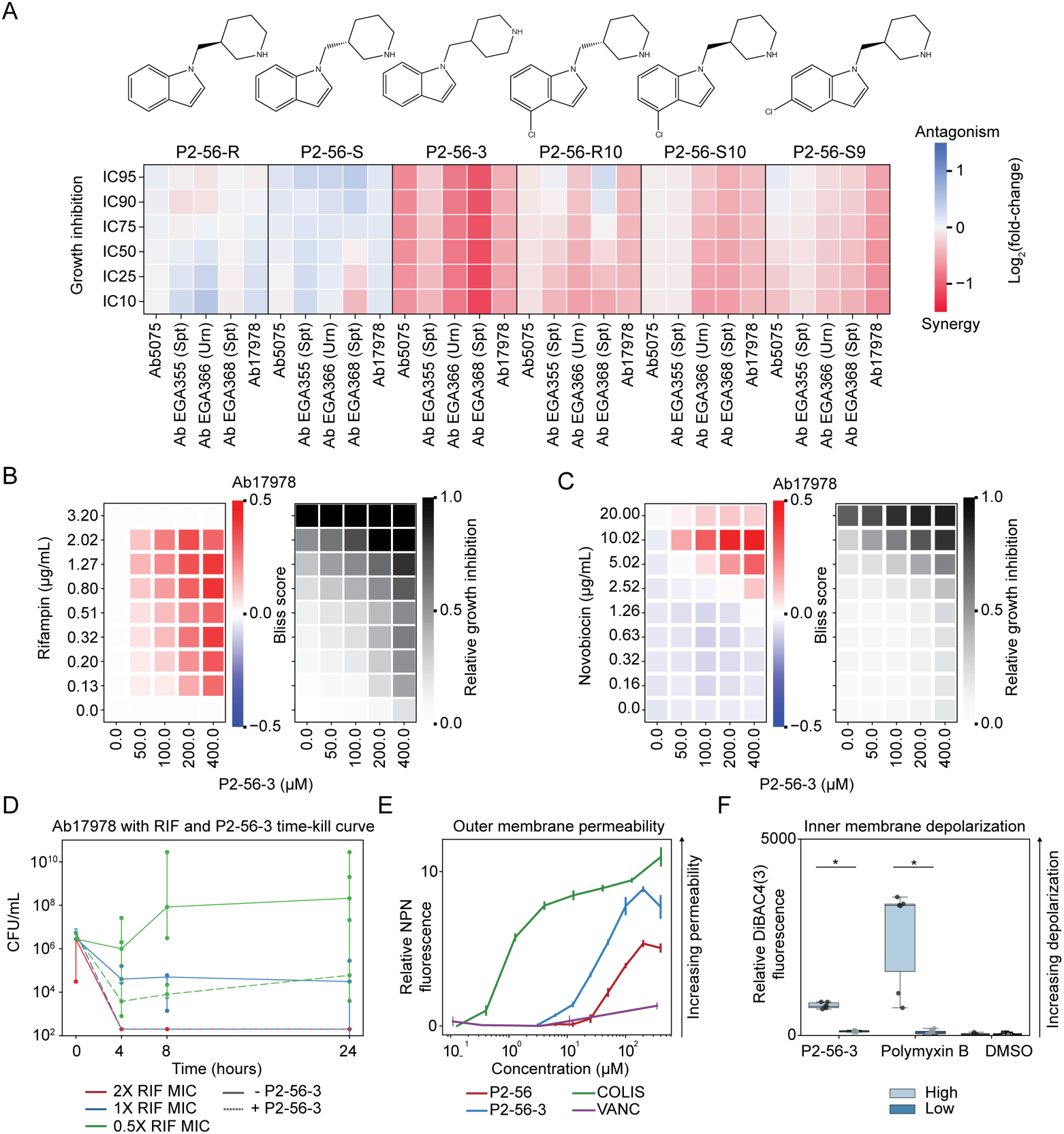
P2-56-3 increases outer membrane permeability. (A) Structural analogs of P2-56 tested in combination with RIF at varying concentrations in 5 strains of *A. baumannii*. Log_2_(fold-change) of the growth inhibition with RIF with 50 µM of compound relative to growth inhibition with RIF only (synergy in red and antagonism in blue). Plate-based checkerboards showing Bliss scores (left) and growth inhibition (right) to evaluate the drug interactions between P2-56-3 and (B) RIF and (C) novobiocin in *A. baumannii* ATCC 17978 (mean of 2 technical replicates and 2 biological replicates). (D) Time-kill curve of *A. baumannii* ATCC 17978 treated with RIF concentrations with and without P2-56-3. Colony forming unit (CFU) concentrations for all three technical replicates represented. Data shown and error bars represent the median ± range. Limit of detection is 200 CFU/mL. (E) Relative NPN fluorescence, as a measure of outer membrane permeability, of *A. baumannii* ATCC 17978 treated with P2-56-3, colistin (positive control), and vancomycin (negative control). Data shown and error bars represent the mean ± SD (3 technical replicates and 1 representative biological replicate). (F) Relative DiBAC4(3) fluorescence, as a measure of inner membrane depolarization, of *A. baumannii* ATCC 17978 treated with P2-56-3, polymyxin B (positive control), and DMSO (solvent control). Data represented as box plots show median and interquartile range. * p ≤0.05 for difference between high and low concentrations. (E, F) P-value, FDR, and/or relative fluorescence calculations: Materials and Methods.

Additionally, we investigated the synergistic effects of P2-56-3 with RIF in other clinically-relevant Gram-negative and Gram-positive pathogens, including *K. pneumoniae*, *P. aeruginosa*, *E. coli, S. aureus*, and *E. faecium*. We observed P2-56-3/RIF synergy in *K. pneumoniae* ATCC 43816 and *E. coli* K-12 MG1655 (Fig. S6A,B); however, we observed little to no synergy with *P. aeruginosa* AR0095 and Gram-positive species, *S. aureus* Newman and *E. faecium* BAA-2317 (Fig. S6C-E). Thus, P2-56-3 exhibited synergistic activity with RIF in all of the tested non-pseudomonal Gram-negative species and none of the Gram-positive species, suggesting P2-56-3 may be acting on the outer membrane.

### P2-56-3/RIF combination treatment kills *A. baumannii* and disrupts its outer membrane

We further characterized the inhibitory effects of the P2-56-3/RIF combination treatment in *A. baumannii* by performing a time-kill assay with multiple concentrations of RIF. As expected, RIF used at 0.5X and 1X its minimum inhibitory concentration (MIC) in *A. baumannii* resulted in no decrease in viability (measured by CFU, colony forming units), demonstrating bacteriostatic activity; whereas, RIF used at 2X its MIC resulted in a sharp decrease in viability, demonstrating strong bactericidal activity (Fig. 3D). Interestingly, RIF used at 0.5X its MIC in combination with P2-56-3 resulted in a slight decrease in viability and RIF used at 1X its MIC with P2-56-3 resulted in a decrease in viability (>3 orders of magnitude at 4 hours), similar to that observed with the 2X MIC RIF only treatment. Thus, we show that P2-56-3 potentiates bactericidal activity of RIF in *A. baumannii*.

Based on P2-56-3/RIF’s synergy observed primarily with Gram-positive-acting antibiotics, selective activity in Gram-negative organisms, and bactericidal activity observed in *A. baumannii*, we hypothesized that P2-56-3 disrupts the outer membrane. To test this hypothesis, we evaluated *A. baumannii*’s outer membrane integrity with and without putative membrane disrupters from the P2-56 series using the N-Phenyl-1-naphthylamine (NPN)-based outer membrane permeability assay (28) (Materials and Methods). Consistent with our previous data, we observed increased relative NPN fluorescence with higher concentrations of P2-56 and P2-56-3, indicating outer membrane disruption allowing for NPN dye entrance and intracellular accumulation (Fig. 3E). Additionally, P2-56-3 treatment resulted in greater outer membrane disruption compared to its parent compound P2-56, consistent with its improved synergistic activity against *A. baumannii* in combination with RIF. Similarly, we observed increased outer membrane permeability with P2-56 and P2-56-3 in *A. baumannii* in a different lysozyme-based permeability assay (29). Since lysozyme typically cannot penetrate the outer membrane of Gram-negative bacteria, additional membrane-disrupting activity is required to induce lysis activity. Thus, the enhanced lysis observed in *A. baumannii* ATCC 17978 treated with lysozyme and P2-56 or P2-56-3 further demonstrates their outer membrane disrupting activity. In the lysozyme assay, greater membrane disrupting activity again occurred with P2-56-3 compared to P2-56 at the same concentration (Fig. S7). To evaluate the specificity of P2-56-3 to the outer membrane, we tested its effect on inner membrane depolarization using the DiBAC3(4) dye uptake assay (30). P2-56-3 treatment produced only minor increases in fluorescence (Fig. 3F) compared to the polymyxin B positive control treatment, suggesting that P2-56-3 does not act strongly on the inner membrane and its activity may be more specific to the outer membrane.

### Essential gene depletions show membrane-compromised *A. baumannii* strains exhibit hypersensitivity to P2-56-3/RIF treatment

Given the minor growth inhibitory effects of P2-56-3 at high concentrations in *A. baumannii* (Fig. 3B), we hypothesized that P2-56-3 could be a weak binder of an essential gene and cause fitness defects in the presence of hypomorphic gene depletions of its pathway or related parallel pathways. To identify genetic determinants of synergy between P2-56-3 and RIF, we conducted a pooled essential gene depletion screen in *A. baumannii* under various drug treatments (Fig. 4A; Dataset S4). We tested the effects of no treatment, 100 µM P2-56-3 only, 0.7 µg/mL RIF only, and 0.7 µg/mL RIF with 100 µM P2-56-3 (P2-56-3/RIF) in the presence of stoichiometric CRISPRi-based gene depletions – also known as hypomorphic gene depletions – in *A. baumannii* ATCC 17978 (31, 32). We looked for hypomorphic gene depletions that induced hypersensitivity to RIF, P2-56-3, and P2-56-3/RIF treatments using the following treatment comparisons: RIF only treatment relative to no treatment, P2-56-3 only treatment relative to no treatment, and P2-56-3/RIF treatment relative to RIF only treatment.

**Figure 4.**
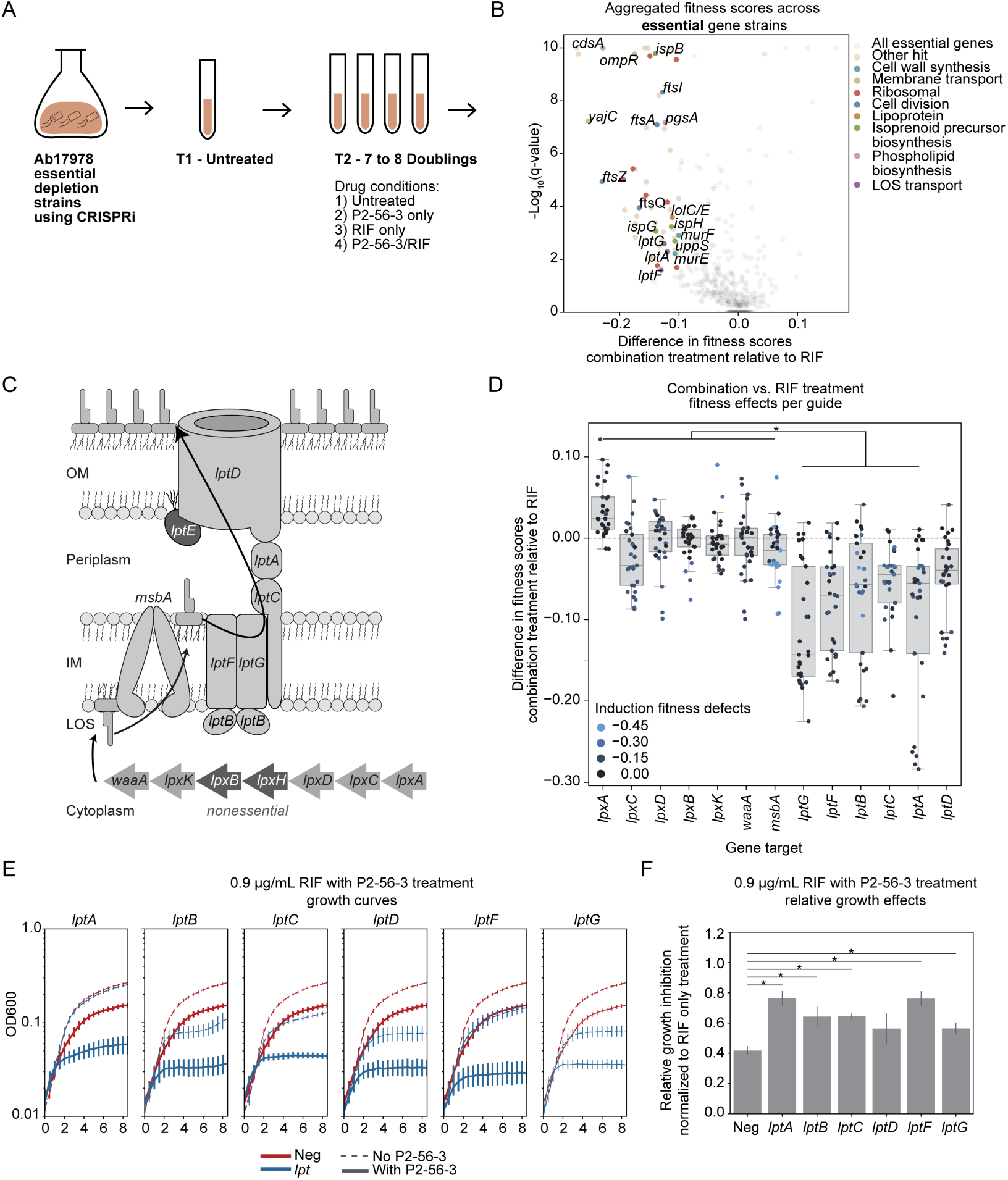
Inhibition of LOS transport in *A. baumannii* enhances fitness defects in the presence of P2-56-3. (A) Schematic of the P2-56-3 and RIF challenge experiments using essential gene depletion strains. Banks of ∼12,000 CRISPRi mutants were challenged with 4 different drug conditions: untreated, 100 µM P2-56-3, 0.7 µg/mL RIF, and the combination of 0.7 µg/mL RIF and 100 µM P2-56-3. (B) Difference in aggregated fitness scores of essential gene depletion strains treated with the combination and RIF only and their significance values represented. Gene depletion strains exhibiting the highest differences in fitness scores labeled (FDR q-value ≤0.05 & fitness difference ≤-0.1). (C) Diagram of LOS synthesis (*lpx)* in the cytoplasm and transport (*lpt*) across the inner and outer membranes (37, 38). (D) Difference in fitness scores between the combination treatment and RIF only for all *lpt* and *lpx* gene depletions. All sgRNA guides targeting each gene represented. Data represented as box plots show median and interquartile range. (E) Growth curves and (F) relative growth inhibition of LOS transport and negative control (Neg) gene depletion strains treated with 0.9 µg/mL RIF only and in combination with P2-56-3 at 8 hours. *: FDR q-value ≤0.05 for comparison between depletion strain and the negative control. (E, F) Data shown and error bars represent the mean ± SEM if shown (4 technical and 2 biological replicates). (B, D, F) P-value and FDR calculations: Materials and Methods.

Consistent with RIF’s mechanism of action, depletions of the RNA polymerase subunit encoded by *rpoD* resulted in strong fitness defects in the presence of RIF relative to no treatment (Fig. S8A,B). We observed that gene depletions associated with envelope biogenesis, particularly in LOS transport, LOS biosynthesis, cell division, and cell wall synthesis resulted in hypersensitivity to RIF only treatment (in agreement with previous work) and RIF/P2-56-3 treatment, implicating these pathways as potentially important to P2-56-3/RIF synergy (Fig. S8A; Fig 4B) (33–35). P2-56-3 only treatment also revealed significant, although modest, fitness defects in LOS transport and cell division gene depletion strains (Fig. S8C). We hypothesize these smaller effect sizes are consistent with the lack of independent inhibitory activity of P2-56-3 observed at this concentration.

Importantly, gene depletions in LOS transport, cell division, and cell wall synthesis exhibited even greater fitness defects with the P2-56-3/RIF treatment relative to RIF only treatment (Fig. 4B). However, LOS biosynthesis gene depletions did not exhibit hypersensitivity with P2-56-3/RIF relative to RIF treatment only despite having exhibited strong fitness defects with RIF only relative to no treatment (Fig. 4B; S8A). To further demonstrate the specificity of these effects to depletions in the LOS transport pathway and not the LOS biosynthesis pathway, we analyzed the individual fitness changes of the LOS biosynthesis and LOS transport depletion strains from the original CRISPRi screen. LOS is synthesized in the cytoplasm and transported to the outer leaflet of the outer membrane through the LOS transport complex (Fig. 4C) (36, 37). We observed all individual LOS transport depletion strains, except *lptD*, had significantly larger fitness defects on average compared to the individual LOS biosynthesis depletion strains (Fig. 4D). These results demonstrate that hypersensitivity to the P2-56-3 and P2-56-3/RIF combination treatment is specific to depletions in the LOS transport pathway (Fig. 4D; S9A). In contrast, RIF alone caused strong fitness defects in both individual LOS biosynthesis and transport depletion strains (Fig. S9B). Thus, the essential gene depletion data showed genes involved in LOS transport, cell division, and cell wall synthesis exacerbate P2-56-3/RIF synergy and likely relate to the molecular mechanisms underlying the interaction between P2-56-3 and RIF.

### LOS transport gene depletions in *A. baumannii* result in P2-56-3 hypersensitivity

Since gene depletions in LOS transport exhibited some of the largest and most significant fitness defects across multiple treatment conditions (P2-56-3/RIF relative to RIF only treatment, P2-56-3 relative to no treatment, and RIF only relative to no treatment), we created *A. baumannii* strains with corresponding gene depletions and tested them in an arrayed format to confirm their hypersensitivity to P2-56-3/RIF. We included a non-targeting guide control strain as our negative control for comparison to ensure that relative fitness defects with treatment were due to the gene depletion and not the CRISPRi induction. We then evaluated the growth of each arrayed gene depletion strain with the following treatment conditions: RIF only, P2-56-3 only, P2-56-3/RIF, and no treatment. Similar to the pooled screen, we observed increased growth inhibitory effects in the LOS transport (*lpt*) gene depletion strains (*lptA*, *lptB, lptC*, *lptF,* and *lptG)* relative to the negative control with *lptA* exhibiting the greatest enhanced growth defect with P2-56-3/RIF relative to RIF only treatment (Fig. 4E,F). Specifically, we observed a nearly two-fold increase in growth inhibition in the *lptA* depletion strain relative to the negative control at 8 hours (76.4 ± 4.5% relative growth inhibition with *lptA* depletion and 41.8 ± 2.6% relative growth inhibition with the negative guide between P2-56-3/RIF and RIF only treatment using 0.9 µg/mL RIF; Fig. 4E,F). Given the minor fitness defects observed under P2-56-3 only treatment in the CRISPRi screen, we additionally tested the isolated single *lpt* depletion strains under P2-56-3 only treatment. The majority of *lpt* strains constructed showed minor growth inhibitory effects, although they were not significant, under P2-56-3 only treatment (Fig. S10A,B). Taken together, we identified LOS transport as an important pathway to the mechanism of synergy between RIF and P2-56-3.

### Nonessential disruption of envelope genes enhances P2-56-3 effects

To further investigate pathways impacting in the synergy between P2-56-3 and RIF, we searched for treatment hypersensitivity in nonessential gene knockout mutants of *A. baumannii* generated using high-density transposon mutagenesis (Fig. 5A; Dataset S5; Materials and Methods) (33, 38). We hypothesized that nonessential gene knockouts resulting in enhanced fitness defects with P2-56-3 only relative to no treatment and with P2-56-3/RIF relative to RIF only treatment would point towards parallel pathways associated with the mechanism of interaction between P2-56-3 and RIF.

**Figure 5.**
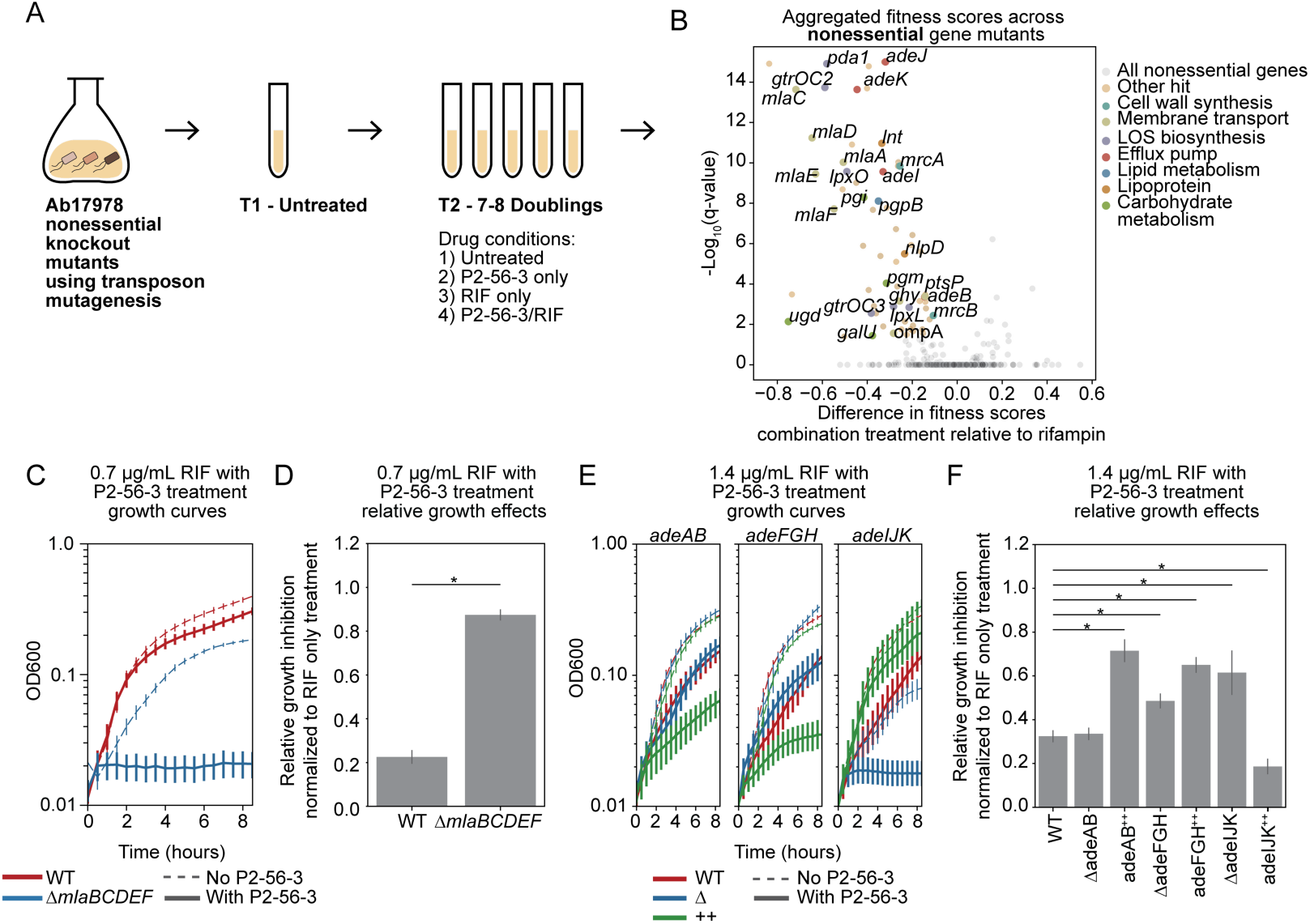
P2-56-3 is more effective in *A. baumannii* strains with compromised outer membranes. (A) Schematic of P2-56-3 and RIF challenge experiments using nonessential gene knockout mutants. Transposon knockout mutants were challenged with 4 different drug conditions: untreated, 100 µM P2-56-3, 0.7 µg/mL RIF, and the combination of 0.7 µg/mL RIF with 100 µM P2-56-3. (B) Difference in aggregated fitness scores of nonessential gene knockout mutants treated with the combination and RIF only and their significance values represented. Gene knockout mutants exhibiting the highest differences in fitness scores labeled (FDR q-value ≤0.05 & fitness difference ≤-0.1). (C) Growth curves and (D) relative growth inhibition at 8 hours of MLA knockout mutant treated with 0.7 µg/mL RIF only and in combination with P2-56-3. *: p-value ≤0.05 for comparison between knockout mutant and WT. (E) Growth curves and (F) relative growth inhibition at 8 hours of *adeAB*, *adeFGH*, and *adeIJK* knockout and hyperexpresser strains treated with 1.4 µg/mL RIF only and in combination with P2-56-3. *: FDR q-value ≤0.05 for comparison between mutant and WT. (C**–**F) Data shown and error bars represent the mean ± SEM if shown (4 technical and 3 biological replicates). (B, D, F) P-value and/or FDR calculations: Materials and Methods.

Similar to the CRISPRi gene depletion results, knockout of genes associated with maintaining envelope integrity, such as cell wall synthesis, resulted in mutants with the most significant fitness defects under P2-56-3/RIF combination relative to RIF only treatment (Fig. 5B). In particular, knocking out genes associated with the maintenance of lipid asymmetry (MLA) protein complex (*mlaA, mlaC, mlaD, mlaE, mlaF*) resulted in hypersensitivity to P2-56-3/RIF relative to RIF only treatment (Fig. 5B). The MLA complex is responsible for maintaining lipid asymmetry between the inner and outer leaflet of the outer membrane (38, 39). Consistent with the gene-aggregated fitness scores from the transposon mutagenesis experiment, individual MLA knockout mutants had minor fitness defects in the presence of RIF only and P2-56-3/RIF caused further sensitization (Fig. S11A). Likewise, a total MLA deletion mutant (Δ*mlaBCDEF*) grown in the presence of 0.7 µg/mL RIF was sensitized by the addition of P2-56-3 (88.9 ± 2.2% relative growth inhibition comparing the P2-56-3/RIF and the RIF only treatment) relative to the wildtype (WT; 22.5 ± 1.8% relative growth inhibition comparing the P2-56-3/RIF and the RIF only treatment) at 8 hours (Fig. 5C,D). We observed similar effects at 0.32 µg/mL RIF (Fig. S11B,C). Given the strong effects of P2-56-3/RIF treatment we observed with the *ΔmlaBCDEF* mutant, we tested the *ΔmlaBCDEF* mutant with a higher dose of P2-56-3. P2-56-3 only treatment resulted in increased growth defects of the *ΔmlaBCDEF* mutant over the WT (Fig. S11D,E) with minimal solvent effects (Fig. S11F). Together with the results from the CRISPRi depletions, we observe knockouts of outer membrane-associated gene products, such as components of the MLA complex, confer P2-56-3/RIF hypersensitivity.

### Overexpression of RND efflux pumps hypersensitize *A. baumannii* to P2-56-3

Previous work demonstrated that overproduction of resistance-nodulation-cell division (RND) efflux pumps in *A. baumannii* sensitized cells to genetically-induced outer membrane disruption (38). RND pump overexpression, a phenotype observed in multidrug-resistant strains, contributes to intrinsic antibiotic resistance and RIF tolerance (41). Moreover, multidrug-resistant clinical isolates of *A. baumannii* exhibit or spontaneously induce RND efflux pump overexpression through mutations in regulating proteins (i.e. mutations in adeS (42, 43), adeL (44), and adeN (42) activate *adeAB*, *adeFGH*, and *adeIJK*). Therefore, we hypothesized that RND efflux pump-overexpressing strains would be more susceptible to P2-56-3 sensitization of RIF relative to wild-type. Thus, we evaluated the effects of P2-56-3 and RIF treatment in individual overexpression (^++^) and deletion (Δ) strains of the RND pumps, *adeAB*, *adeFGH*, and *adeIJK* (Materials & Methods) (38).

With combination treatment, we observed decreased growth of the *adeIJK* deletion (Δ*adeIJK*) mutant relative to the WT (Fig. 5E,F; S12A,B). We expected this decreased growth, given the previously observed fitness defects of the transposon knockout mutants (Fig. 5B) and the intrinsic resistance to RIF known to be associated with *adeIJK*. Unlike the Δ*adeIJK* mutant, deletion of *adeAB* (Δ*adeAB)* and *adeFGH* (Δ*adeFGH*) did not significantly inhibit growth (Fig. 5E,F; S12A,B). We observed significantly increased susceptibility to 1.4 µg/mL RIF with P2-56-3 in the *adeAB* overexpression strain (*adeAB^++^*; 71.4 ± 4.9% relative growth inhibition between P2-56-3/RIF and RIF only treatment; activating *adeS*) and *adeFGH* overexpression strain (*adeFGH^++^*; 65.0 ± 3.4% relative growth inhibition between P2-56-3/RIF and RIF only treatment; activating *adeL*) when compared to WT (32.4 ± 2.5% relative growth inhibition between P2-56-3/RIF and RIF only treatment) at 8 hours (Fig. 5E,F). We observed this trend of increased growth inhibitory effects at 0.7 µg/mL RIF as well (Fig. S12A,B). In contrast, we did not observe increased growth inhibition in the *adeIJK* overexpression (*adeIJK*^++^; inactivating *adeN*) mutant. Notably, P2-56-3 exhibited greater effectiveness as a single-agent treatment in the *adeAB^++^*, *adeFGH^++^* and *adeIJK^++^* strains compared to the WT (Fig. S12C,D) where the DMSO solvent minimally affected growth of these strains (Fig. S12E). Overall, we demonstrated that P2-56-3 exhibited greater effectiveness as a potentiator for RIF and a growth inhibitory single-agent drug in the presence of RND pump hyperexpression, indicating that pump overproduction paradoxically increases sensitivity to this compound.

## Discussion

In this work, we expanded the capacity of DropArray to assay over 1.3 million unique strain-antibiotic-compound combinations encompassing up to 6 different strains of the Gram-negative ESKAPE pathogens, 22 antibiotics, and over 30,000 natural, semi-synthetic, and synthetic compounds. In total, we called 3,784 antibiotic-compound combination hits across 2,388 unique library compounds. Overall, antagonistic antibiotic-compound interactions dominated synergistic interactions; however, we observed that synergistic effects with existing antibiotics still occurred more frequently than independent compound effects, highlighting the opportunity to identify new compound synergies in phenotypic screens exploring large combinatorial chemical spaces. Our phenotypic screening effort discovered novel synergies in the highly resistant *A. baumannii* LAC-4, *K. pneumoniae* AR0087, and *P. aeruginosa* AR0095 strains, and a number of instances of stronger antibiotic synergy in more resistant strains, demonstrating the promise of phenotypic combination screening to nominate new therapeutic hypotheses relevant in the setting of highly antibiotic resistant infections of high priority Gram-negative ESKAPE pathogens.

DropArray enabled the discovery of strong synergistic effects of P2-56 with 8 antibiotics across 5 strains in the primary screening data where the strongest synergy occurred with Gram-positive-specific antibiotics against strains of *A. baumannii* and *K. pneumoniae.* We synthesized a structural analog, P2-56-3, which potentiated RIF against *A. baumannii* and *K. pneumoniae* more than the parent compound. We suspect that P2-56-3 did not synergize with RIF in *P. aeruginosa* given this organism’s widely recognized high levels of intrinsic antibiotic resistance due to especially low outer membrane permeability (45).

Using plate-based phenotypic assays, we demonstrated that P2-56-3 compromised the outer membrane of Gram-negative ESKAPE pathogens. To identify genes affecting P2-56-3/RIF synergy, we depleted essential genes and looked for hypersensitization of *A. baumannii* to the P2-56-3/RIF treatment. Our results demonstrated that depletion of proteins involved in cell envelope homeostasis – particularly LOS transport genes – hypersensitized bacteria to P2-56-3/RIF over RIF only treatment and P2-56-3 only treatment relative to untreated. RIF is a hydrophobic molecule that normally has difficulty penetrating the outer membrane of Gram-negative bacteria (13). These data suggest that P2-56-3 could be weakly affecting LOS transport or a parallel pathway to compromise outer membrane integrity. Previous screens of *A. baumannii* with RIF treatment also showed decreased fitness associated with mutants defective in division, cell wall synthesis, membrane transport, LOS biosynthesis and transport (33–35). Additionally, the recent discovery of a new antibiotic class binding LptB_2_FG highlighted LOS transport as an important and effective target for single-agent antibiotic development against *A. baumannii* (46, 47). In our study, we highlight LOS transport genes, including *lptF* and *lptG*, as genetic determinants for the activity of P2-56-3 and its potentiation of RIF efficacy, expanding the scope of candidate targets and further highlighting the therapeutic potential of inhibiting this complex. LOS, in particular, is important for maintaining outer membrane integrity. The negative charge of LOS at the outer membrane disfavors the penetration of hydrophobic molecules, contributing to antibiotic resistance and thus preventing RIF accumulation (48, 49). Thus, we hypothesize that the membrane permeabilizing activity of P2-56-3 allows for increased uptake of hydrophobic molecules such as RIF.

Further supporting the CRISPRi knockdown screen, we identified that high-density transposon mutagenesis of nonessential genes involved in maintaining outer membrane homeostasis, such as the MLA genes, resulted in hypersensitivity under P2-56-3 only relative to no treatment and P2-56-3/RIF relative to RIF only treatment. While CRISPRi screening uncovered a set of genes whose disruption exacerbated P2-56-3 treatment effects and these provide support for an outer membrane integrity centered mechanism of action for P2-56-3 and antibiotic interaction, none of the associated gene products have been confirmed as molecular targets of P2-56-3. Some approaches for identification of direct targets may be challenging to practice, for example the minor growth inhibitory effects of P2-56-3 treatment alone are expected to make it more difficult to raise resistant mutants that can be mapped to putative direct targets.

While overexpression of the *adeAB* or *adeFGH* RND efflux pumps increases resistance to a number of clinically useful antibiotics, overexpression of these drug pumps hypersensitized *A. baumannii* to treatment with P2-56-3 alone and its combination with RIF. Multidrug-resistant clinical strains predominantly overexpress at least one RND efflux pump (41). Recent work demonstrated RND efflux pump overexpression causes hypersensitivity to *A. baumannii* strains with outer membrane defects (38). We suspect that the combined stress of pump overexpression and P2-56-3 could cause increased outer membrane disruption compared to strains lacking pump overexpression, thus facilitating antibiotic action. In contrast, we observed no hypersensitization with a mutant overexpressing *adeIJK*, an RND efflux pump that can transport RIF as a substrate, indicating that *adeIJK* hyperactivity may counteract potential loss of envelope integrity by reducing stress from RIF activity. By this model, strains overexpressing *adeIJK* may also be hypersensitized to P2-56-3 in combination with antibiotics that are not substrates of this pump.

In addition to the proven ability of our platform to discover diverse synergistic interactions between compounds and clinically useful antibiotics, there are other advantages of DropArray. The deep antibiotic interaction profile from the primary screen provides information useful for prioritizing initial hits. For example, we observed that P2-56 amplified the activity of several Gram-positive-acting antibiotics, consistent with a mechanism based on outer membrane disruption and growth inhibition of Gram-negative pathogens. The inclusion of multiple species and strains further proves an initial indication of the activity spectrum. While the work reported here demonstrates the scalability of DropArray for antibiotic potentiator discovery, we need to deploy phenotypic combination screening more broadly. This broader deployment is necessary to generate a continuous stream of translatable drug combinations capable of addressing the growing threat of multidrug-resistant bacterial pathogens. We think that subsets of chemical space with enriched bioactivity (9), ADMET properties (50), or structural features by empirical evidence or computational prediction represent a rational place to start (51, 52). The datasets generated in this study may serve as training data for future antibiotics discovery efforts employing machine learning methods or improve our understanding of chemical properties associated with antibiotic potentiators (53–55).

In summary, we demonstrated DropArray’s capability to identify compounds that enhance and expand the activity of existing antibiotics. Using this strategy, we re-identified known and discovered new antibiotic-compound combinations that result in enhanced antibiotic activity against the high-priority, multidrug-resistant Gram-negative ESKAPE pathogens. We discovered the potentiator activity of compound P2-56-3, likely attributable to disruption of the outer membrane of Gram-negative strains and explored its synergistic effects with RIF against RND efflux pump-overexpressing *A. baumannii* strains, which are associated with multidrug resistance in the clinic. This study shows that phenotypic DropArray screening can be used to find synergistic drug interactions with activity against clinically-relevant strains with a broad range of resistance profiles.

## Materials and Methods

### Resource availability

#### Lead contact

Further information and requests for resources and reagents should be directed to and will be fulfilled by the lead contacts, Paul C. Blainey and Ralph R. Isberg.

#### Data and code availability

DropArray strain-antibiotic-compound combination (individual and summed Bliss scores of combinations and independent effects of compounds), transposon mutagenesis nonessential genes screen (aggregated fitness scores), and CRISPRi partial depletion essential genes screen (individual and aggregated fitness scores) are available as supplement datasets on the Harvard Dataverse (https://dataverse.harvard.edu/privateurl.xhtml?token=23f475d0-c94e-4ad7-aa2b-2cac89c6196e). Any additional data in the manuscript are available upon request. Code used for DropArray (https://github.com/megantse/abxcombos-droparray), CRISPRi (https://github.com/megantse/abxcombos-crispri), and Tn-seq analysis (https://github.com/megantse/abxcombos-tnseq) are uploaded to GitHub.

## Method details

### Strains and growth conditions

GFP-expressing strains of *A. baumannii* ATCC 17978 (25 µg/mL tetracycline selection and 50 µM IPTG inducible), *A. baumannii* LAC-4 (50 µg/mL tetracycline selection and 50 µM IPTG inducible), *K. pneumoniae* ATCC 43816 (10 µg/mL chloramphenicol selection and constitutive), *K. pneumoniae* AR0087 (10 µg/mL chloramphenicol selection and constitutive), *P. aeruginosa* PAO1 (50 µg/mL gentamicin selection and constitutive), and *P. aeruginosa* AR0095 (50 µg/mL gentamicin selection and constitutive) were used in the Otava library screen. The same strains, except for a constitutive GFP-expressing strain of *A. baumannii* ATCC 17978 (100 µg/mL carbenicillin selection and constitutive) instead of the previously mentioned inducible strain, were used in all other screens. For checkerboard assays, *E. coli* K-12 MG1655, *S. aureus* Newman, and *E. faecium* ATCC BAA-2317 were additionally used. All assays were performed in Cation-Adjusted Mueller Hinton II Broth (CAMHB, BD Difco #L007475) at 37°C and end point reads for growth in plate assays were taken at 8 hours unless otherwise indicated. Details on strains, plasmids, and oligonucleotides used are listed (Table S4, S5).

### Compound sourcing and analog generation

The following compound libraries were obtained: Otava (Otava, acquired from the Broad Institute Center for the Development of Therapeutics), Natural Products (Microsource Discovery Systems), Charles River (Charles River Laboratories), Diversity (Diversity-oriented synthesis compound collection, Broad Institute) (18, 19), and CtoD (Complex-to-Diversity compound collection, provided by Paul Hergenrother’s lab) (20, 21). Compounds with reproduced synergy were reordered and resynthesized (VITA Biotech). Analogs of hit compounds were generated by a contract research organization (VIVA Biotech). The synthesis protocol of each tested P2-56 analog is available upon request.

### DropArray Screens: antibiotic and compound preparation; bacterial preparation; and droplet emulsification, pooling, loading, and imaging

We previously described the DropArray method for antibiotic combinations screening and is summarized below (17). CAMHB was mixed with a unique ratio of 3 fluorescent dyes, Alexa Fluor 555, 594, and 647 (Invitrogen #A33080, #A33082, #A33084) at a final concentration of 20 µM for each unique assay input. Antibiotics dissolved in DMSO or water with 0.01% Triton-X100 were prepared at 2-3 concentrations in CAMHB with their designated barcoding dye (Table S6; Dataset S1) using the Tecan D300e Digital Dispenser. Compound libraries including Otava and Natural Products were prepared at a final concentration of 100 µM in CAMHB with their designated barcoding dye. Other compound libraries including Charles River, Diversity, and CtoD were prepared at a final concentration of 50 µM in CAMHB with their designated barcoding dye.

Single colonies were selected from LB agar plates with the corresponding antibiotic selection and grown overnight in 5 mL of CAMHB at 37°C with shaking at 250 rpm and antibiotic selection. Overnight cultures were diluted in fresh media and mixed with the prepared antibiotic solutions prior to emulsification with a final assay OD_600_ of 0.02. Arrayed emulsifications for each assay input were prepared using a BioRad QX200 droplet generator using 7500 Engineered Oil (3M Novec) with 2% 008-fluorosurfactant (RAN Biotechnologies) as the oil phase. Droplets were pooled, mixed, loaded into the discoveryChip, and sealed using an optically clear PCR film (Applied Biosystems) pre-wetted with 7500 Engineered Oil.

The discoveryChips were imaged with a fluorescent microscope with a Nikon 1x Plan Achromat Microscope objective. For the Otava and Natural Products libraries, we used a Nikon Ti-E fluorescence microscope equipped with the Lumencor Sola light engine, the Hamamatsu Orca Flash 4.0 camera, and the following filter cubes: Alexa Fluor 555: Semrock SpGold-B; Alexa Fluor 594: Semrock 3FF03-575/25–25 + FF01-615/24–25; Alexa Fluor 647: Semrock LF635-B; and GFP: Semrock GFP-1828A. For the other libraries, we used a Nikon Ti-2 fluorescence microscope equipped with the Lumencor Sola light engine, the Iris 9 camera, and the same filter cubes. After the discoveryChips were loaded and sealed, we imaged prior to droplet merging in the 4 channels corresponding to the filter cubes listed previously. Droplets were then merged using a corona treater (Electro-Technic Products #Model BD-20). At the designated timepoints per species (*A. baumannii* and *K. pneumoniae* 6.5 to 7 hours and *P. aeruginosa* 12 hours) after static incubation at 37°C, we imaged the merged discoveryChips across the 4 channels again to obtain bacterial growth information using GFP expression.

### Data analysis

A Snakemake analysis pipeline was developed for the integration of DropArray image analysis and antibiotic potentiation scoring with previously developed custom Python scripts (17). Briefly, in the premerge images, the pairwise combination of droplets in each microwell was decoded using the ratio of the fluorescent barcodes. The positioning of each microwell was then mapped to the merged images with the GFP readout. Antibiotic and novel compound interactions were measured using Bliss scores. Bliss scores are calculated as the difference between the observed combination growth inhibitory effect (*E*_*ac*_) and the expected combination growth inhibitory effects (*Ea* + *Ec* − *Ea* ∗ *Ec*), where *E*_*a*_ indicates growth inhibition effects by the antibiotic and *E*_*c*_ indicates growth inhibition effects by the compound. To calculate a scalar statistic for each antibiotic-compound combination, we summed the individual Bliss scores to get Bliss sum scores. To estimate the standard error, we bootstrapped the Bliss score (1000 iterations) and calculated standard deviation. Then, we computed a test statistic by dividing the Bliss sum score by the summed standard error. The test statistic was then modeled with a t-distribution that was fit to the negative controls to obtain p-values. We calculated false discovery rate (FDR) q-values using the Benjamini-Hochberg procedure (statsmodel 0.11.0). Combination hits were called using a Bliss sum score ≥0.3 and FDR q-value ≤0.05. Independent single-agent hits were called using a growth inhibition ≥30% and FDR q-value ≤0.05. FDR q-values were calculated on a batch basis per library.

Chip quality was measured using Z-prime, which was determined based on the GFP signal from the combinations representing the maximum and minimum signal. For the maximum signal, we used the median signal (*m*_+_) of the bacteria only paired with media combinations and the corresponding standard error (*s*_+_). For the minimum signal, we used the median signal (*m*_−_) of the media paired with media combinations and the corresponding standard error (*s*_−_). The following equation was used: 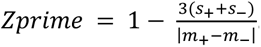. We filtered out chips with Z-primes ≤0.2 for our hit calling procedure.

### DropArray hit validation assays

To validate combination hits from the primary screen, compounds were reordered and tested at more than one concentration in a checkerboard format on DropArray. Antibiotic concentrations were selected to cover the full range of growth. For Otava and Natural Products, specific combinations were retested in full checkerboard formats with 7–8 compound concentrations in DropArray. For the Charles River set, compounds were selected for reduced checkerboard formats with 2 compound concentrations, 10 and 50 µM, in DropArray. For the Diversity and Hergenrother set, compounds were selected for reduced checkerboard formats with 2 compound concentrations, 25 and 50 µM, in DropArray. DropArray chips were prepared and imaged as previously described. For Otava and Natural Products, Bliss sum scores in the validation assays were calculated across 100 µM compound concentrations and the 7–8 antibiotic concentrations tested. For Charles River, Diversity, and Hergenrother libraries, Bliss sum scores in the validation assays were calculated across 50 µM compound concentrations and the 5 antibiotic concentrations tested. Compounds with Bliss sum scores ≥0.3 with at least 1 strain-antibiotic pair, calculated across the set of antibiotic concentrations tested with the compound concentration used in the primary screen (50 or 100 µM), were called as reproduced hits. For single-agent hits, compounds were tested across a range of concentrations up to 400 µM in DropArray against all screening strains. Compounds were indicated as reproduced for independent activity if growth inhibitory effects were larger than 30% at 50 µM for at least one strain.

### Plate-based checkerboard assays

For plate-based evaluation of compounds, similar or matching conditions to those tested on the DropArray were performed in microtiter plates. 384W plates were prepared with the relevant drug treatments, dispensed using the Tecan D300E Digital Dispenser. For P2-56 analog testing, overnight cultures were prepared as described above and then diluted 1:500 in 3 mL of CAMHB and grew up to OD_600_ 0.2–0.5. Prior to starting the assay, the cultures were then diluted to OD_600_ 0.001, and 50 μl culture was added to 384W plates. For other checkerboard assays, overnight cultures were prepared as described above and then diluted to OD_600_ 0.1. Prior to starting the assay, the cultures were then diluted and added to 384W plates for a 80 µL total culture volume and a final assay OD_600_ of 0.003. Plates were grown overnight at 37°C with shaking and OD_600_ was measured using a microplate reader (Spectramax M3, Spectramax M5, Agilent Biotek Cytation 5, or Agilent Biotek Synergy HT). At least 2 biological replicates were performed for each combination and strain.

### Time-kill curves

Overnights of *A. baumannii* ATCC 17978 were grown from single colonies in CAMHB at 37°C on a shaker. Overnights were diluted to OD_600_ 0.1 and grown up at 37°C on a shaker until approximately OD_600_ 0.2-0.5. Cultures were diluted to OD_600_ 0.003 prior to starting the assay and drug treatment was added in 1 mL total cultures in a 24W deep well plate. Treated and untreated cultures were grown at 37°C on a shaker. At 0, 4, 8, and 24 hours, 20 µL was removed for colony forming unit (CFU) plating and counting. To estimate CFU count, we made 10-fold serial dilutions in PBS starting with neat, plated 5 µL on LB agar, and incubated plates overnight at room temperature for manual colony counting the next day. If no colonies were observed at the neat concentration, the CFU/mL were plotted at the limit of detection (<200 CFU/mL). 3 biological replicates were performed for each condition: 2, 4, and 8 µg/mL RIF with or without 400 µM P2-56-3.

### Drug challenge of CRISPRi depletion strain pools

CRISPRi depletion pools were generated as previously described in *A. baumannii* ATCC 17978 (32). To briefly summarize, the CRISPRi library contained 12,000 guides targeting 406 essential genes, including one perfect match and 23 mismatched guides per gene and included 2,000 scrambled, non-targeting negative controls (32). The mismatched guides were designed with 1 or 2 mismatches at varying distances from the PAM site to allow for varying fitness defects and thus stoichiometric gene depletions. Frozen aliquots of *A. baumannii* ATCC 17978 hypomorphic strains containing the plasmid bank were thawed and diluted to OD_600_ 0.1. Cultures were grown until OD_600_ was approximately 0.2 and then normalized to OD_600_ 0.003 at the first time point (*t*1) in 3 mL cultures in CAMHB with induction using 50 ng/mL aTc. The following treatment conditions were applied to the induced cultures using 1.5 mg/mL RIF and 100 mM P2-56-3 stocks in DMSO: 1) Untreated (DMSO control), 2) 100 µM P2-56-3, 3) 0.7 µg/mL RIF, and 4) 0.7 µg/mL RIF with 100 µM P2-56-3. Cultures were approximately grown to OD_600_ 0.5–1 at the second time point (*t*2) at 37°C on a roller. Samples were stored at -20°C until DNA extraction and sequencing. The combination treatment conditions selected resulted in approximately 30% growth rate inhibition. 6 biological replicates of the libraries were tested across all treatment conditions.

### CRISPRi Illumina library preparation and sequencing

Plasmid DNA containing sgRNA was extracted using the QIAprep Spin Miniprep Kit (Qiagen #27104). sgRNAs were amplified in 50 µL reactions with 100 ng plasmid DNA using For2_sgRNA_Pool and Rev_sgRNA_Pool and Q5 polymerase (New England Biolabs #M0491L). sgRNA were further amplified with Nextera indexes using a mix of N5x index primers and N7x index primers in a 10 µL reaction with 1 µL of the previously amplified sgRNAs. Barcoded samples were then quantified on a 2% agarose gel with a known DNA ladder standard and then pooled at 50 ng each. After purification, pooled samples were submitted for sequencing using single-end 100 bp reads with an Illumina HiSeq 2500 at the Tufts University Genomics Core Facility.

### CRISPRi data analysis

Sequencing reads were adapter trimmed using cutadapt (v3.4) and mapped to sgRNAs using a custom python script (count_guides.py) (57). Mapped reads were then used to calculate individual sgRNA depletion strain fitnesses as described. First, we normalized sgRNA reads by the total mapped reads per sample by summing the column of all mapped reads per sample. sgRNAs with cumulative read numbers less than 20 at *t*1 were excluded from the analysis for that condition. Raw zeros were transformed to read count of 1 to calculate fitness. The fitness W was calculated based on an individual mutant vs. population-wide expansion between *t*1 and *t*2 with the following equation: 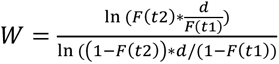 where *F*(*t*2) and *F*(*t*1) are the normalized mutant frequency at *t*2 and *t*1 and *d* is the population expansion. These fitness scores were further normalized to a set of neutral/negative controls. To obtain an aggregated score, the median of the fitnesses across all guides and replicates used. For each drug treatment, the difference in fitness per gene was calculated relative to the untreated and deemed significant with a FDR q-value ≤0.05. We calculated FDR q-values using an unpaired, two-tailed t-test with FDR control by the two-stage step-up method of Benjamini, Krieger, and Yekutieli (SciPy 1.6.2).

### Drug challenge of transposon mutant pools

Transposon mutant banks were generated as previously described (34) and challenged with drugs. Briefly, frozen banks of *A. baumannii* ATCC 17978-derived transposon mutants (5,000-20,000 random mutants per bank with a total of 21 banks) were thawed and diluted to OD_600_ 0.1. Cultures were grown until approximately OD_600_ 0.2 and then normalized to OD_600_ 0.003 at the first time point (*t*1) in 5 mL cultures in CAMHB. The following treatment conditions were applied to each bank using 1.5 mg/mL RIF and 100 mM P2-56-3 stocks in DMSO: 1) Untreated (DMSO control), 100 µM P2-56-3, 3) 0.7 µg/mL RIF, and 4) 0.7 µg/mL RIF with 100 µM P2-56-3. Cultures were grown to approximately OD_600_ 0.5–1 at the second time point (*t*2) at 37°C on a roller. Samples were stored at -20°C until DNA extraction and sequencing. The combination treatment conditions selected resulted in approximately 30% growth rate inhibition.

### Tn-seq Illumina library preparation and sequencing

Genomic DNA was extracted using a DNeasy Blood & Tissue Kit (Qiagen #69504). Transposon-adjacent DNA was prepared for Illumina sequencing using a modified Nextera DNA library prep method as previously described (58). Briefly, 50 ng genomic DNA was tagmented in a 10 µL reaction mixture. Transposon-adjacent DNA was then amplified using primers olj638 for libraries and Nextera 2A-R. Samples were further amplified using nested indexed primers (left mariner-specific indexing primers and right indexing primers). Samples were run on a 2% agarose gel loaded with a known NEB DNA ladder and quantified based on the 250–600 bp region. Quantified samples were then pooled using 50–70 ng per sample and purified. After the pools were reconditioned using adapter-specific primers P1 and P2 and purified, the pools were submitted for sequencing using single-end 50 bp reads with an Illumina HiSeq 2500 at the Tufts University Genomics Core Facility.

### Tn-seq data analysis

Sequencing reads were adapter trimmed using fastx_clipper, quality filtered using fastq_quality_filter, and mapped to the *A. baumannii* ATCC 17978 genome (annotations acquired from ATCC website) using bowtie. Mapped reads were then used to calculate individual transposon mutant fitnesses using custom scripts previously described (33, 57, 58). The fitness W was calculated based on an individual mutant vs. population-wide expansion between *t*1 and *t*2 with the following equation: 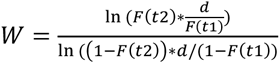 where *F*(*t*2) and *F*(*t*1) are the normalized mutant frequency at *t*2 and *t*1 and *d* is the population expansion. For each treatment condition, gene-level fitness was calculated by averaging the fitness of all transposon mutants across all mutant pools having insertions in the first 90% of the gene and normalized to selected neutral genes. For each drug treatment, the difference in fitness per gene was calculated relative to the untreated and deemed significant with a FDR q-value ≤0.05. We calculated FDR q-values using an unpaired, two-tailed t-test with FDR control by the two-stage step-up method of Benjamini, Krieger, and Yekutieli (SciPy 1.6.2).

### Molecular cloning and isolation of knockout mutants

Cloning inserts were amplified using oligonucleotide primers (Integrated DNA Technologies) and cloned into pJB4648 prior to recombination or gene expression. Gene deletions were constructed using three-way ligation of ∼750 bp flanking homology arms with pJB4648. These deletion constructs were electroporated in *E. coli* DH5α λpir and selected using 10 µg/mL gentamicin. Plasmids were isolated using QIAprep Spin Miniprep Kit (Qiagen #27104) and the regions of interest were confirmed using Sanger sequencing through Genewiz. *A. baumannii* ATCC 17978 was electroporated with deletion constructs and allelic exchange mutants were isolated via homologous recombination and two selection steps. Transformants were selected on 10 µg/mL gentamicin, single colony purified with no selection, counter selected on 10% sucrose twice, and then single colony purified with no selection. All steps with sucrose were carried out at room temperature for 2 days or 30°C overnight. All other steps were carried out at 37°C overnight. Isolation of the deletion mutants was verified by colony PCR. Details on plasmids and oligonucleotides used for generation of knockout strains are listed (Table S4, S5).

### Molecular cloning and isolation of depletion strains

We acquired an *A. baumannii* ATCC 17978 with an aTc inducible dCas9 gene inserted at the chromosomal *att*Tn-7 site downstream of *glmS* locus (32). sgRNA inserts were designed to be 24 nucleotides with TAGT (top) and AAAC (bottom) sticky ends. Inserts were phosphorylated and annealed, prior to ligation into the BsaI-HF2 digested pWH1266. These sgRNA constructs were then transformed into *E. coli* DH5α, isolated using QIAprep Spin Miniprep Kit (Qiagen #27104), and verified with Sanger sequencing (Genewiz). Confirmed sgRNA constructs were electroporated into *A. baumannii* ATCC 17978 with an inducible dCas9 and selected on 50 µg/mL carbenicillin overnight at 37°C. Details on plasmids and oligonucleotides used for generation of knockdown strains are listed (Table S4, S5).

### Verification of fitness defects observed in CRISPRi and Tn-seq screens

Knockout mutants of *A. baumannii* ATCC 17978 were grown up overnight in 37°C in 1 mL CAMHB. Overnight cultures were diluted to OD_600_ 0.1 and then grown until OD_600_ 0.2–0.5. Upon starting the assay, knockout mutants were added to 384W plates under RIF and/or 200 µM P2-56-3 treatment with a final starting OD_600_ 0.003. The corresponding negative control was the wildtype strain.

Depletion strains of *A. baumannii* ATCC 17978 dCas9 were grown overnight in 37°C in 1 mL CAMHB with 50 µg/mL carbenicillin. Prior to starting the assay, overnights were diluted to OD_600_ 0.1 and then grown until OD_600_ 0.4–0.6. Upon starting the assay, depletion strains were added to 384W plates under RIF and/or 200 µM P2-56-3 treatment with a final starting OD_600_ 0.003. Depending on the depletion effect, 50 or 100 ng/mL aTc was added. The corresponding negative control was a strain containing a plasmid with a scrambled, non-targeting guide that caused no fitness defects.

Relative growth was calculated as the OD_600_ at 8 hours for each condition relative to the untreated control. For the compound only treatment, the relative growth inhibition normalized to the untreated was calculated by the following formula: *relative growth inhibition* = 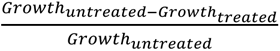. For the combination compared to RIF only treatment, the relative growth inhibition normalized to the RIF only treatment was calculated by the following formula: *relative growth inhibition* = 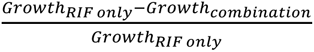. For compound only treatment, we used an unpaired, two-tailed t-test to check if growth in the mutant strains was lower or higher than the negative control strain at the same concentrations. For the combination treatment, we used an unpaired, two-tailed t-test to compare the differences in combination treatment to RIF only treatment relative to the negative control strain. When the number of tests were greater than 5, we used q-values with FDR control by Benjamini-Hochberg to call significance with q-values less than 0.05 deemed as significant. Otherwise, we called p-values less than 0.05 as significant. At least 2 biological replicates and 4 technical replicates were performed for each condition.

### NPN outer membrane assay

The NPN (Sigma-Aldrich #104043) dye was used to assay outer membrane permeability. 1 mM stock solution of NPN was prepared in acetone (Thermo Scientific #22928) and diluted to 40 µM in 5 mM HEPES (Gibco #15630080). Overnight cultures of *A. baumannii* ATCC 17978 were inoculated at 37°C in 10 mL LB. Overnight cultures were diluted 1:50 in LB until OD_600_ 0.4–0.6. Cultures were spun down and resuspended in 5 mM HEPES on ice. 100 µL of bacterial suspension was mixed with 100 µL of 20 µM NPN in 5 mM HEPES with or without drug for testing conditions in a 96W black, clear bottom plate. Vancomycin and colistin were used as controls, negative and positive respectively, because of their known mechanisms. Fluorescence was measured at 355 nm excitation and 405 emission using a microplate reader (Spectramax M5) after 1 hour incubation at room temperature. Normalized scores were calculated based on the final timepoint. The relative fluorescence values were calculated as a ratio of background-corrected (subtracted by the average values in the absence of NPN) fluorescence values of the bacterial suspension and of the buffer, respectively and solvent subtracted. 3 technical replicates were performed.

### Lysozyme outer membrane assay

For the lysozyme assay, overnights of *A. baumannii* 17978 were inoculated at 37°C in LB. Overnights were backdiluted 1:50 in LB until 0.5 OD_600_. Cultures were spun down and resuspended in 5 mM HEPES on ice. 2 mL microcentrifuge tubes were prepared in advance with 95 µL of 5 mM HEPES with or without drug for testing conditions. Vancomycin and colistin were used as controls, negative and positive respectively, because of their known mechanisms. 5 µL of 10 mg/mL lysozyme stock was added to each tube. 900 µL of the cultures in 5 mM HEPES was added to each tube and mixed thoroughly through tube inversion. The mixture was left for 15 minutes at room temperature to allow debris to settle. OD_600_ was then measured on a 96W clear bottom plate using a microplate reader (Spectramax M5). Percent lysis was calculated by dividing OD_600_ values by the untreated condition. 3 technical replicates were performed. Statistical significance was tested using one-way ANOVA with Tukey’s multiple comparisons across all conditions.

### DiBAC4(3) membrane depolarization assay

*A. baumannii* ATCC 17978 was inoculated overnight from single colonies in 10 mL LB. 10 mg/µL DiBAC4(3) (Invitrogen #B438) in DMSO were prepared and stored at -20°C. Overnight cultures were diluted to OD_600_ 0.01 and grown to OD_600_ ∼0.3 at 37°C. Cultures were normalized to OD_600_ 0.3 and then 3 mL cultures were grown at 37°C under the following drug treatments: P2-56-3 (100 and 400 µM), DMSO (0.1 and 0.4%), and polymyxin B (0.5 and 2.5 µg/mL). Polymyxin B was used as a positive control. After 3 hours, OD_600_ was measured and 2 mL of culture was removed and spun down. Final concentration of 10 µg/mL DiBAC4(3) was added and incubated at 37°C in the dark. After 5 minutes, the cultures were spun down and resuspended in 1X PBS. 100 µL was plated and fluorescence was measured at 460 nm excitation and 518 nm emission using a 96W black, clear bottom plate. The relative fluorescence values were calculated using the measured background subtracted RFUs normalized to the background subtracted OD_600_ and subtracted from the untreated control. 3 biological replicates and 2 technical replicates were performed. We calculated p-values, using an unpaired, two-tailed t-test between the high and low treatments.

## Acknowledgments

We thank members of the Isberg, Mecsas, Aldridge, Hung, and Blainey labs, in particular Rachel Ende, Rebecca Carlson, Anna Le, and Robert Majovski, for valuable feedback and discussion. We thank members of the Center for the Development of Therapeutics at the Broad Institute for chemistry support. We thank Paul Hergenrother’s lab for providing the CtoD library. This work was completed as part of the Center for Innovation to Transform Antibiotic Discovery at the Broad Institute and MIT (CITADel) with support from NIH NIAID Grant U19AI42780, as well as funds from the Broad Institute of MIT and Harvard, the MIT Deshpande Center Innovation Award, the Merkin Institute for Transformative Technologies in Healthcare, the Koch Institute for Integrative Cancer Research at MIT, and the Defense Advanced Research Projects Agency. M.W.T. and M.Z. were supported by the National Science Foundation Graduate Research Fellowship under Grant No. 1745302. J.C. was supported by the Natural Sciences and Engineering Research Council of Canada Postgraduate Scholarship – Doctoral (NSERC PGS-D No. 567809 - 2022). J.M. received support from NIH Grant R01AI169789 and Y.M. received support from NIH Grant T32AI007422. R.R.I. received support from NIH grants U19AI158076, 1U01AI124302-01, and R21AI128328-02.

