## Supplemental Information for "Massively parallel combination screen reveals small molecule sensitization of antibiotic-resistant Gram-negative ESKAPE pathogens"

1. Department of Biological Engineering, Massachusetts Institute of Technology, Cambridge, MA 02139
2. Broad Institute of MIT and Harvard, Cambridge, MA 02142
3. Department of Molecular Biology and Microbiology, Tufts University School of Medicine, & Stuart B. Levy Center for Integrated Management of Antimicrobial Resistance, Boston, Massachusetts, 02111
4. Microbiology Graduate Program, Massachusetts Institute of Technology, Cambridge, MA 02139
5. Tri-Institutional Program in Computational Biology and Medicine, New York, New York, 10065
6. Wellcome Sanger Institute, Hinxton, Saffron Walden CB10 1RQ, United Kingdom
7. Department of Molecular Biology and Center for Computational and Integrative Biology, Massachusetts General Hospital, Boston, MA, 02114
8. Tango Therapeutics, Boston, MA, USA 02215
9. Department of Chemistry, Massachusetts Institute of Technology, Cambridge, MA 02139
10. Department of Genetics, Harvard Medical School, Boston, MA 02115
11. Department of Biomedical Engineering, Tufts University, Medford, MA, 02155
12. These authors contributed equally.
13. These authors contributed equally.
14. These authors are co-corresponding and contributed equally.

**Corresponding authors**:

Paul C. Blainey
Ralph R. Isberg

**This PDF file includes:**

Supporting text, including supporting methods
Figures S1 to S13
Tables S1 to S6
Legends for Datasets S1 to S7
SI References

**Other supporting materials for this manuscript include the following:**

Datasets S1 to S7

### Supporting Information Text

#### Natural products and semi-synthetic compounds are more likely to cause growth inhibitory effects compared to synthetic compounds

We evaluated the trends observed across the growth inhibitory compounds that were called hits (Supplemental Methods; Dataset S6). Of note, the Natural Products library comprised natural products while the CtoD library comprised semisynthetic compounds. In addition, the Otava, Charles River, and Diversity libraries comprised synthetic compounds. When evaluating the compound growth inhibitory hit set across each library across strains (Fig. S13A), we observed higher hit rates with the natural products and semi-synthetic compounds. Consistent with previous antibiotics discovery efforts, we expected this higher hit rate for natural products given that synthetic compound screening historically has yielded low hit rates (<0.001%) and natural products are known to exhibit enriched bioactivity (1, 2).

We evaluated the differences in physicochemical properties between the single-agent hit and the non-hit sets (Supplemental Methods; Dataset S6). In addition to a lower pKa, we observed that the number of hetero-rings, amides, aromatic nitrogens, and acidic oxygens were significantly lower. The number of H-bond donors was significantly higher among the single-agent hits. Previous studies have indicated that antibacterials effective against Gram-negative bacterial strains typically have lower logD, more H-bond donors, and higher polar surface areas compared to drugs from the Comprehensive Medicinal Chemistry database (3, 4). These differences were likely due to the stringent requirements for bacterial cell entry. Previous studies have shown that compounds with the following properties enrich compound uptake in Gram-negative bacteria: positive charges (primary amine-containing in particular), low globularity, and a low number of rotatable bonds (5). In particular, these observations were obtained by studying accumulation in *E. coli*. We likely did not observe overlap in these properties with our hit set because our screen studied growth inhibition (rather than accumulation) in a distinct set of Gram-negative species (*A. baumannii*, *K. pneumoniae*, and *P. aeruginosa*).

**Compounds are more likely to be antagonistic than synergistic with known antibiotics**

We evaluated the trends across the synergistic and antagonistic interaction hit combinations (Supplemental Methods; Dataset S7). We generally observed higher rates of synergistic interactions among the natural products or semi-synthetic compounds compared to fully synthetic compounds (Fig. S13B). Among all the unique strain-antibiotic-compound combinations, the synergistic hit rate was 0.18% (2,152/1,209,363) and the antagonistic hit rate was 2.50% (30,232/1,209,363). We observed antagonism more frequently than synergy for every strain, antibiotic, and library in our study (Fig. S13B,C). Previous studies have observed higher rates of antagonism over synergy across pairwise drug interactions of antibiotics with diverse mechanisms in Gram-negative bacteria (6–8). Even when combining antibiotics with compounds lacking significant independent growth inhibitory effects, the trend of observing higher antagonism over synergy remained apparent.

We evaluated the differences in physicochemical properties between the compounds involved in the synergistic combination hit and the non-hit sets (Supplemental Methods; Dataset S7). In addition to lower molecular weight, pKa, van der Waals volume, van der Waals surface area, total surface area, and globularity (calculated with singular value decomposition), we observed a decreased number of H-bond acceptors, rotatable bonds, aromatic nitrogens, hetero-rings, amides, symmetric atoms, electronegative atoms, ring closures, sp3-atoms, small rings, saturated rings, amines, aromatic rings, basic nitrogens, alkyl-amines, non-aromatic rings, and aromatic amines among the hit molecules. In addition to higher globularity (calculated with van der Waals properties), relative polar surface area, and topological polar surface area, we observed a higher number of acidic oxygens, non-aromatic rings, carbo-rings, stereocenters, and H-bond donors within the hit molecules. We obtained our antibiotic-compound combinations dataset across a large panel of antibiotics with diverse mechanisms. Although no other similar datasets of antibiotic-compound combinations exist, we highlight a previous study focused on discovering outer membrane-targeting agents that resulted in synergy with Gram-positive-acting antibiotics in Gram-negative bacteria (9). This study observed lower logD, logS, and topological polar surface area and higher pKa, number of basic nitrogens, positive charges, sp3 hybridization, and molecular flexibility. We did not observe overlaps in physicochemical properties with the Klobucar et al. study, likely due to differences in the assays performed that resulted in different hit chemical matter. Klobucar et al. focused their assay on detecting outer membrane-targeting agents; however, we used an unbiased growth-based phenotypic assay and likely found antibiotic-compound combination hits with different mechanisms of synergy.

We evaluated the differences in physicochemical properties between the compounds involved in the antagonistic combination hit and the non-hit sets (Supplemental Methods; Dataset S7). In addition to lower acidic pKa, logD, molecular weight, topological polar surface area, total surface area, van der Waals volume, and van der Waals surface area, we observed a lower number of rotatable bonds, H-bond acceptors, electronegative atoms, amides, aromatic nitrogens, hetero-rings, symmetric atoms, aromatic rings, ring closures, small rings, and saturated rings in the hit set. In addition to a higher basic pKa, logS, and globularity (calculated with van der Waals properties), we observed a higher number of H-bond donors, non-aromatic rings, acidic oxygens, carbo-rings, and sp3-atoms among the antagonistic hit set.

We observed a high overlap of chemical properties between the antagonistic and synergistic hit set, likely due to the 61% (813/1,337) overlap of the synergistic hit compounds with the antagonistic set. We expected the high overlap between the synergistic and antagonistic hit sets given that a single antibiotic can have both synergistic and antagonistic effects when tested in combination with several other antibiotics.

### SI Methods

#### Antibiotic inhibitory concentration determination

To determine the inhibitory concentration (IC) of each antibiotic across each strain, we used data from plate-based $OD_{600}$ antibiotic curves. To determine the IC90, we used the minimum antibiotic concentration that resulted in at least 90% growth inhibition at 8 hours. If no values across the tested range exceeded 90% growth inhibition, we used 2X the maximum tested concentration as the approximate IC90 (with maximum values of 512 µg/mL).

#### Bootstrapping analysis for replication determination

To determine the appropriate level of replication for high quality DropArray chips, we performed a power analysis. We generated DropArray chips with high levels of replicates of media only and bacteria only inputs. Using this dataset, we calculated estimated Z-primes by determining median and standard error of the media only and bacteria only GFP values at different replication levels using bootstrapping (1000 iterations), which we repeated 3 times. We then looked for the minimum replication level in which the majority of strains had Z-primes ≥0.2.

#### Chemical representation of the compounds

To generate the t-Distributed Stochastic Neighbor Embedding (t-SNE), we used a previously described method to quantify the chemical relationship between the compounds tested in our study (10). To briefly summarize, we calculated the Morgan fingerprints (RDKit 2020.09.1) using a radius of 2 and 2048-bit fingerprint vectors. Then, we used t-SNE (MulticoreTSNE 0.1) with the Jaccard distance metric to reduce the 2048 dimensions to 2. Since the Jaccard distance is associated with Tanimoto similarity, which is a metric used to evaluate the similarity between chemical structures, the t-SNE is an appropriation representation of the chemical space explored.

#### Fractional inhibitory concentration determination

The fractional inhibitory concentration (FIC) is another test of synergy, in which synergy is defined as FIC ≤0.5. The following equation was used to calculate FIC: $FIC = x/MIC_A + y/MIC_C$ where x is the concentration of antibiotic and y is the concentration of the compound in a combination treatment where growth inhibition exceeded 90%. The MIC values were calculated at 90% growth inhibition after curve fitting the individual effects. We fit the data using the curve_fit function (SciPy 1.6.2) and the Hill function. ICs that were unable to be calculated were selected to be 2X the highest tested concentration. In most cases, we note that the compound had no inhibitory effects. When only 2 concentrations of compound were tested, as was in the case of the Charles River, Diversity, and CtoD compounds, we calculated a slightly different metric, which was the fold change (FC) score at 50 µM: $FC = z/MIC_A$ where z is the concentration of antibiotic where growth inhibition ranged between 50-60% growth inhibition in combination with 50 µM compound. The MIC value was calculated at 50% growth inhibition after curve fitting the antibiotic only effects. We note this was done for hits in checkerboards with dose ranges that did not exceed 90% growth inhibition. We classified the combination as synergistic if the minimum FIC or FC in the matrix was ≤0.5.

**Chemical properties analysis**

Using the SMILES of each compound, a set of chemical descriptors was calculated using properties exported from CDD Vault and DataWarrior (version 5.5.0). After selecting certain criteria for the hit set, we then compared the chemical properties between the compounds that passed the criteria to the non-hit compounds. If properties could not be calculated for certain compounds, those compounds were omitted from the analysis on a per property basis. P-values were calculated using the Mann-Whitney U test and FDR q-values were determined using the Benjamini-Hochberg correction. We called significantly different properties using FDR q-value ≤0.05. This analysis was repeated on a per strain, antibiotic, and/or library basis. For individual compounds, we used the following criteria for hit calling: relative growth inhibition ≥30% and FDR q-value ≤0.05. Compounds that met these criteria for at least 1 strain were deemed a single-agent hit. For combinations, we used the following criteria for hit calling: Bliss sum score ≥0.3 and FDR q-value ≤0.05 for synergy and Bliss sum score ≤-0.3 and FDR q-value ≤0.05 for antagonism. Compounds that met these criteria for at least 1 strain-antibiotic pair were deemed either a synergistic or antagonistic hit.

### SI Figures and Tables

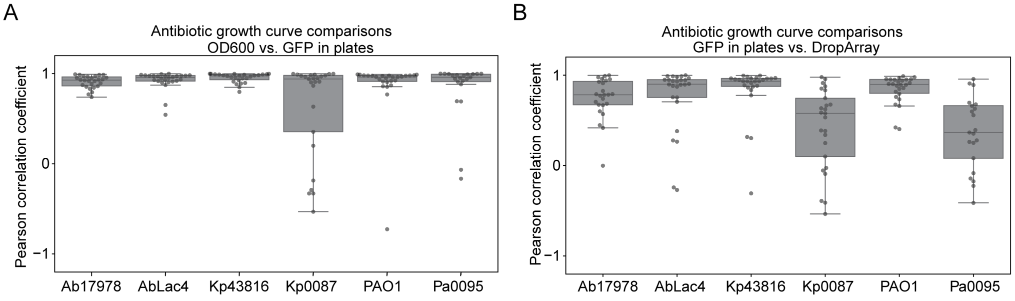

**Fig. S1. Growth inhibition across various readouts, assays, and strains show correlation.** Each point represents a Pearson correlation coefficient calculated between (A) $OD_{600}$ and GFP fluorescence in plates and (B) GFP fluorescence in plates and DropArray chips for selected strain-antibiotic pairs. (A, B) Data represented as box plots show median and interquartile range.

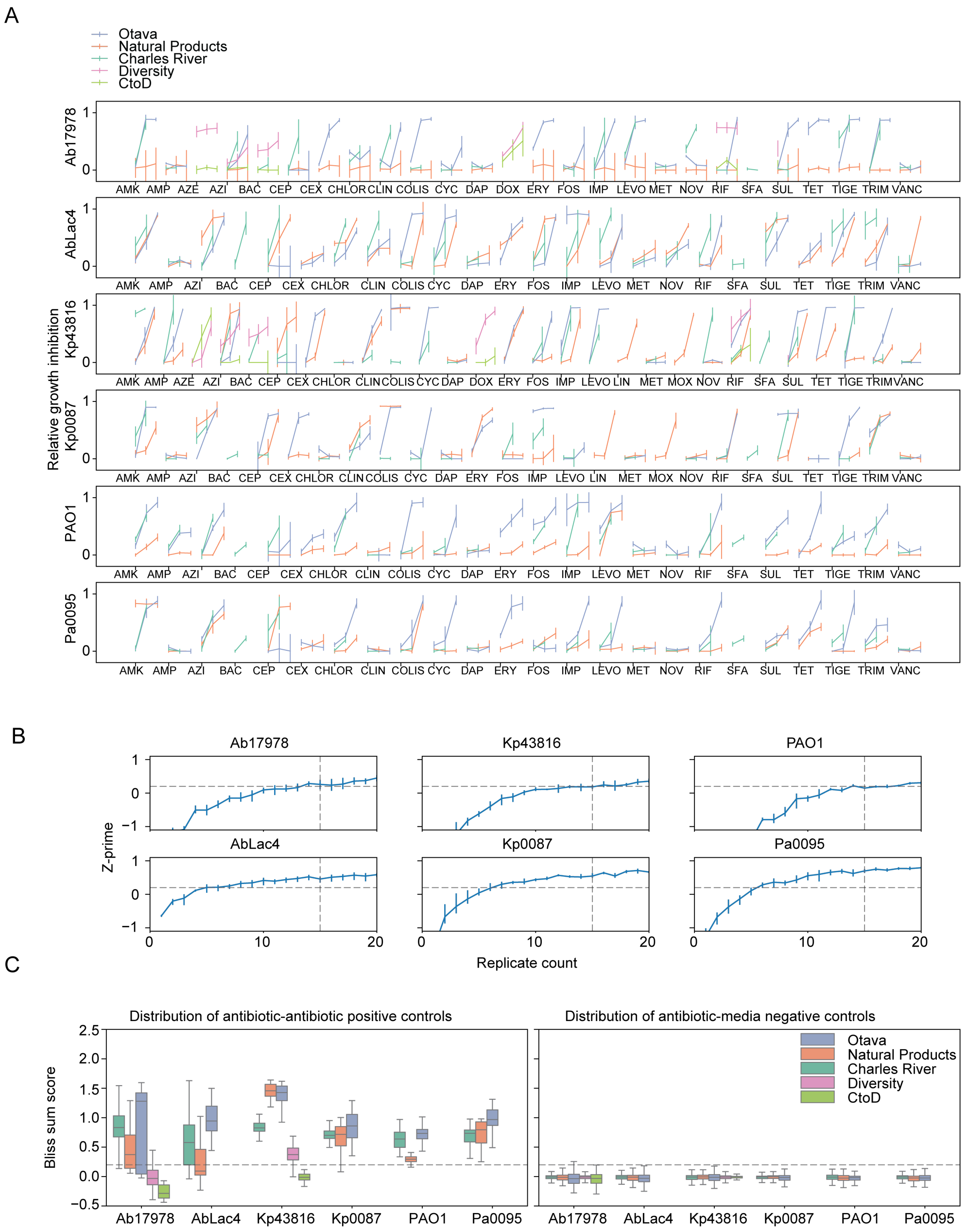

**Fig. S2. Antibiotic growth curves, controls, and replication levels in the primary screen**. (A) (A) Antibiotic concentrations and their relative growth inhibitory effects used for each compound library screen across all strains. For each chip screened that fell above the Z-prime cutoff of 0.2, we determined the median of each antibiotic concentration's growth inhibitory effect. For each strain and antibiotic, data represent the median ± MAD aggregated across all chips. (B) We performed a power analysis for each strain by calculating Z-prime across different replication levels from

DropArray data. We calculated Z-primes using the bacteria only and media only combinations as our positive and negative assay controls. Data shown and error bars represent the mean ± SD across 3 replicates (Materials and Methods). Dotted horizontal line represents the Z-prime cutoff used to quality control filter chips from the primary screen and the dotted vertical line represents the median sampling level observed across all screened chips. (C) We calculated the Bliss sum scores for each antibiotic-antibiotic combination used as a positive synergy control (left) and each antibiotic-media combination used as a negative synergy control (right) per chip across compound libraries. Dotted horizontal line represents the Bliss sum score cutoff used to call antibiotic-compound synergy hits. Data represented as box plots show median and interquartile range.

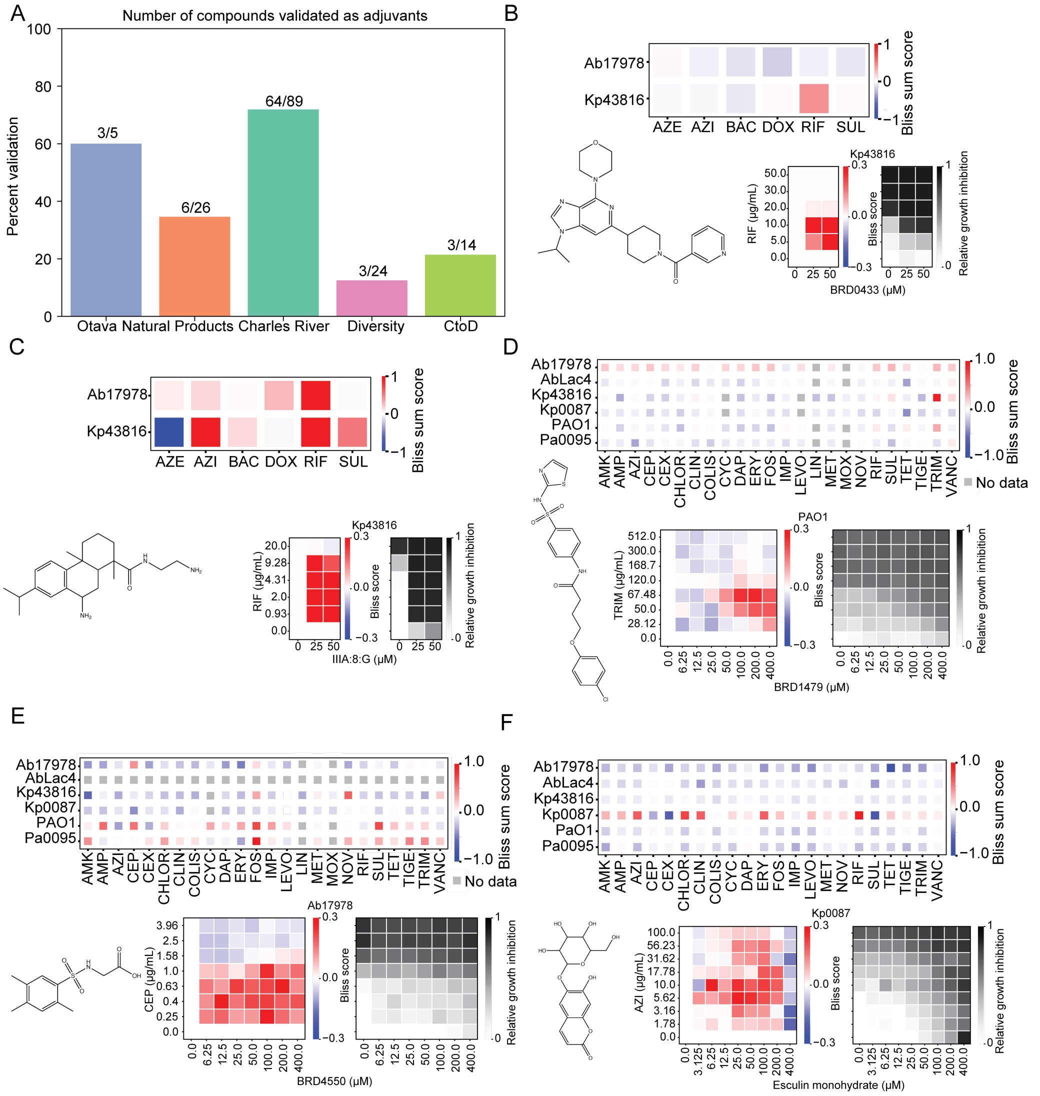

**Fig. S3. Synergistic effects reproduce in checkerboard format of selected antibiotic-compound pairs.** (A) Number of compounds re-tested and validated after DropArray checkerboard assays as potentiators per library. We used the following abbreviations for each strain: *A. baumannii* ATCC 17978 (Ab17978), *A. baumannii* LAC-4 (AbLac4), *K. pneumoniae* ATCC 43816 (Kp43816), *K. pneumoniae* AR0087 (Kp0087), *P. aeruginosa* PAO1 (PAO1), and *P. aeruginosa* AR0095 (Pa0095). The chemical structure, antibiotic interaction profile from the primary screen (synergistic in blue, antagonistic in red, and no data in gray), and DropArray-based checkerboards showing Bliss scores (left) and growth inhibition (right) for selected compound hits: (B) BRD0433 from the Diversity library with rifampin (RIF) against *K. pneumoniae* ATCC 43816, (C) IIIA:8:G from the CtoD library with RIF against *K. pneumoniae* ATCC 43816, (D) BRD1479 from the Otava library with trimethoprim (TRIM) against *P. aeruginosa* PAO1, (E) BRD4550 from the Otava library with cefepime (CEP) against *A. baumannii* ATCC 17978, and (F) esculin monohydrate from the Natural Products library with azithromycin (AZI) against *K. pneumoniae* ATCC 48316.

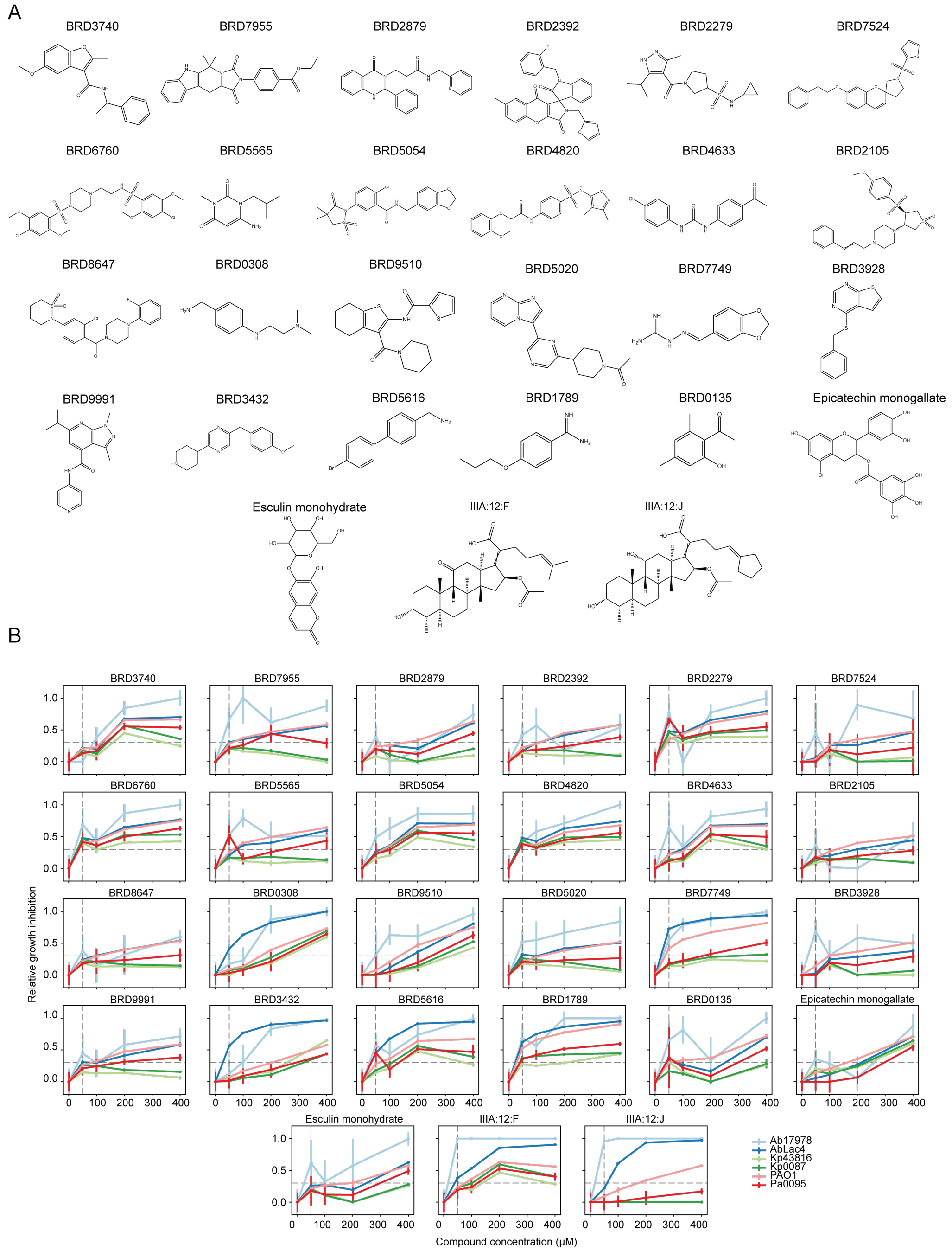

**Fig. S4. Single-agent hits from the primary screen show growth inhibitory effects.** (A) Chemical structures of compounds that validated as a single-agent hit with at least one strain. (B) The corresponding growth inhibition curves for each validated single-agent compound hit for all strains shown across 8 concentrations. The dotted horizontal line represents the compound concentration used in the primary screen and the dotted vertical line represents the relative growth inhibition cutoff used to define single-agent hits. We used the following abbreviations for each strain: *A. baumannii* ATCC 17978 (Ab17978), *A. baumannii* LAC-4 (AbLac4), *K. pneumoniae* ATCC 43816 (Kp43816), *K. pneumoniae* AR0087 (Kp0087), *P. aeruginosa* PAO1 (PAO1), and *P. aeruginosa* AR0095 (Pa0095). Data shown and error bars represent the median ± SE from DropArray.

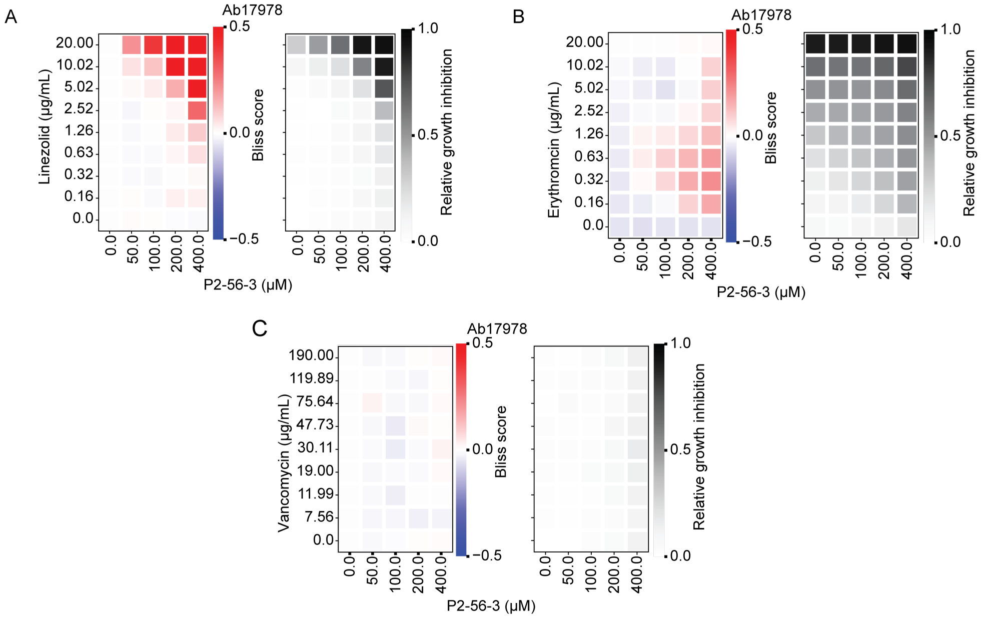

**Fig. S5. P2-56-3 shows synergistic interactions with many Gram-positive-acting antibiotics against *A. baumannii*.** Plate-based checkerboards show Bliss scores (left) and growth inhibition (right) to evaluate drug interactions between P2-56-3 and Gram-positive-acting antibiotics (A) linezolid, (B) erythromycin, and (C) vancomycin with *A. baumannii* ATCC 17978. (A–C) Data represent the mean (2 technical replicates and 2 biological replicates).

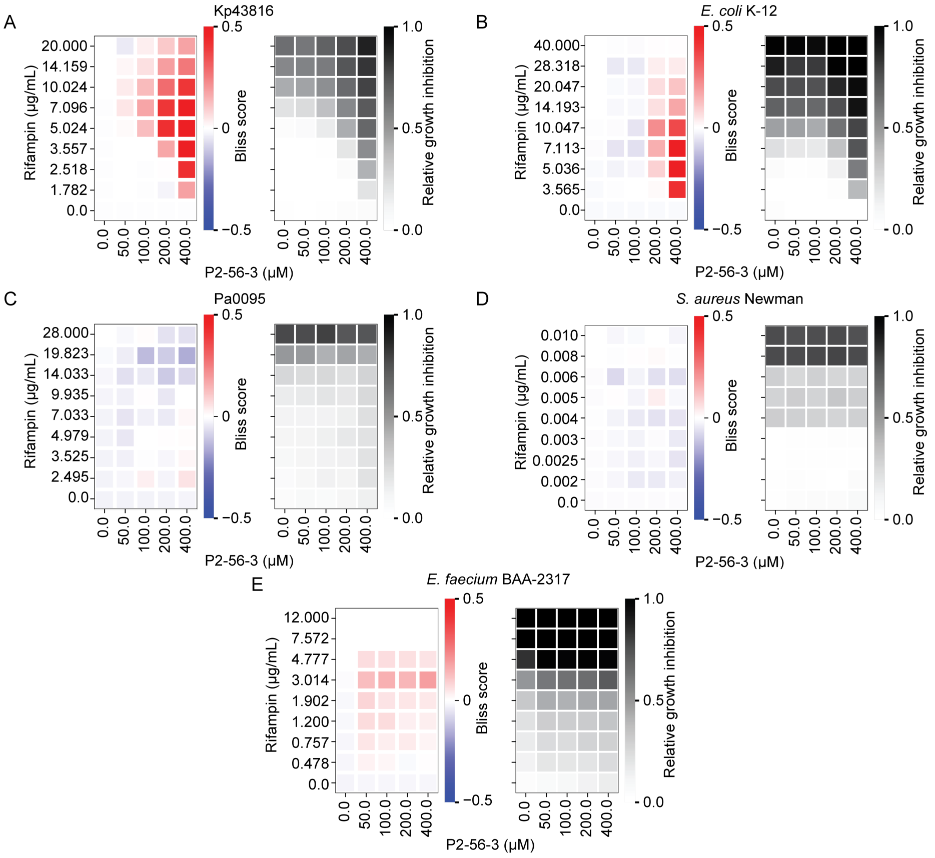

**Fig. S6. Checkerboards between RIF and P2-56-3 show synergistic effects primarily with Gram-negative species.** Plate-based checkerboards show Bliss scores (left) and growth inhibition (right) to evaluate drug interactions between RIF and P2-56-3 in (A) *K. pneumoniae* ATCC 43816 (Kp43816) and (B) E. coli K-12. Little to no interaction was observed between RIF and P2-56-3 in (C) *P. aeruginosa AR0095* (Pa0095), (D) *S. aureus* Newman, and (E) *E. faecium* BAA2317. (A–E) Data represent the mean (2 technical replicates and 2 biological replicates).

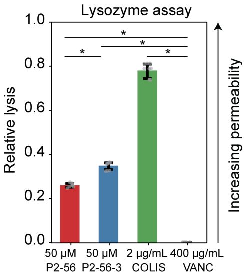

**Fig. S7. Phenotypic assays demonstrate that P2-56-3 permeabilizes the outer membrane.** Lysozyme assay shows increased outer membrane permeability in the presence of P2-56-3, and greater signal over the parent compound. Colistin (positive) and vancomycin (negative) controls. * $p \leq 0.05$. Data shown and error bars represent the mean ± SEM (3 technical replicates and 2 biological replicates). P-value calculations: Materials and Methods.

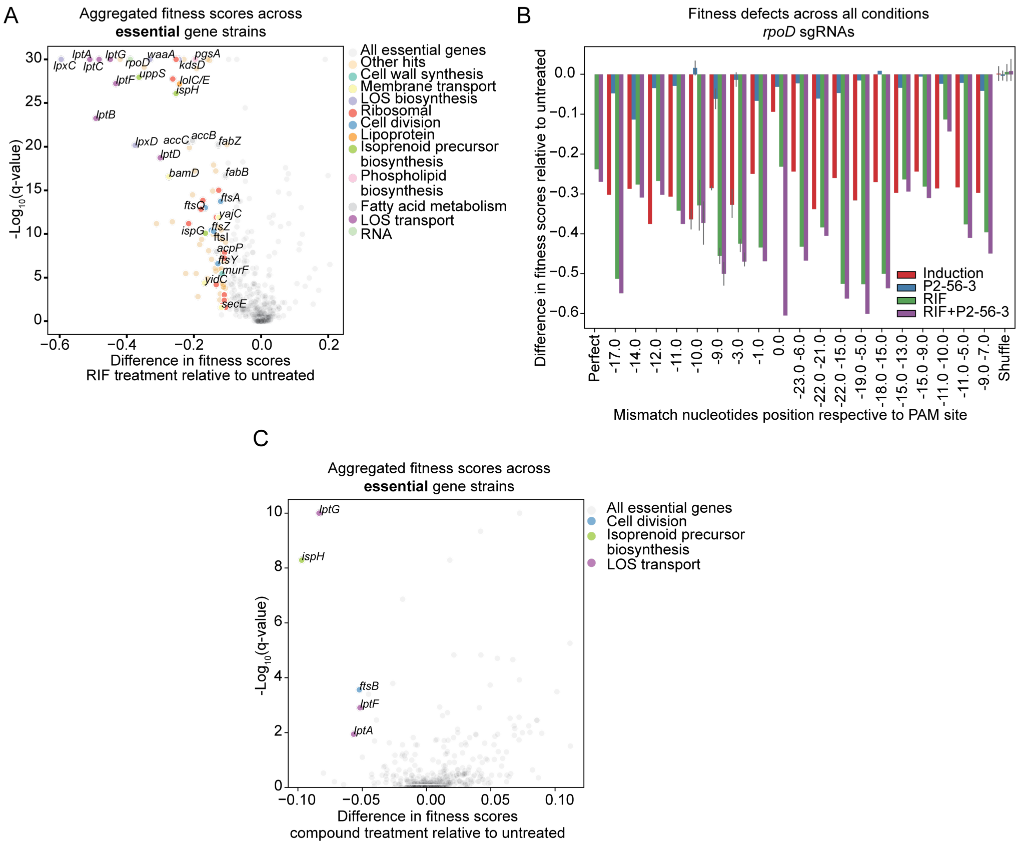

**Fig. S8. Depletions of envelope-associated essential genes reveal similar fitness defects in RIF only and combination treatments.** (A) Difference in aggregated fitness scores of essential gene depletion strains treated with the RIF only relative to the untreated condition and their significance values represented. Gene depletion strains exhibiting the highest differences in fitness scores labeled (FDR q-value ≤0.05 & fitness difference ≤-0.1). (B) Fitness differences induced across all treatment conditions overlaid across the various guides and their mismatch positions relative to the PAM site for *rpoD*. Data and error bars represent the mean ± SD. (C) Difference in aggregated fitness scores of essential gene depletion strains treated with P2-56-3 only relative to the untreated condition and their significance represented. Gene depletion strains exhibiting the highest differences in fitness scores labeled (FDR q-value ≤0.05 & fitness difference ≤-0.05). (A, C) P-value and FDR calculations: Materials and Methods.

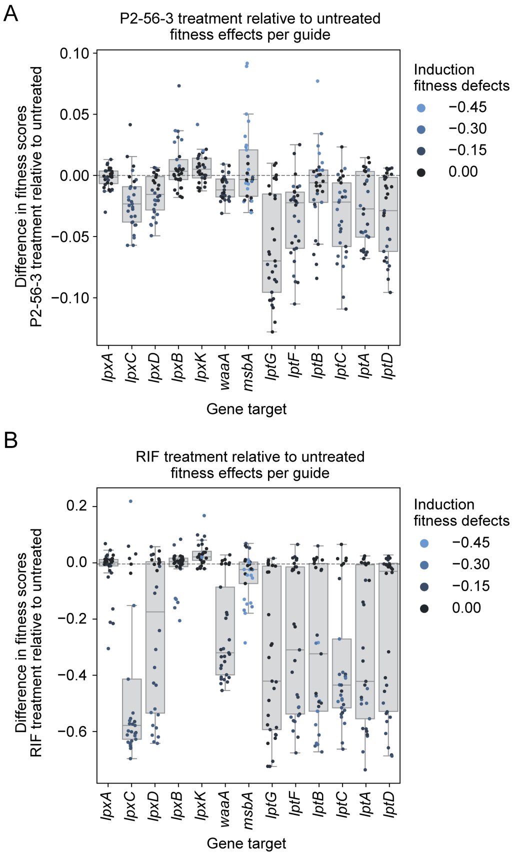

**Fig. S9. Comparison of fitness defects between treatment conditions of individual depletion strains with gene depletions in LOS biogenesis.** Fitness defects from (A) P2-56-3 only treatment and (B) RIF only treatment relative to no treatment for LOS synthesis (*lpx*) and transport (*lpt*) individual gene depletion strains contrasted. (A, B) Data represented as box plots show median and interquartile range.

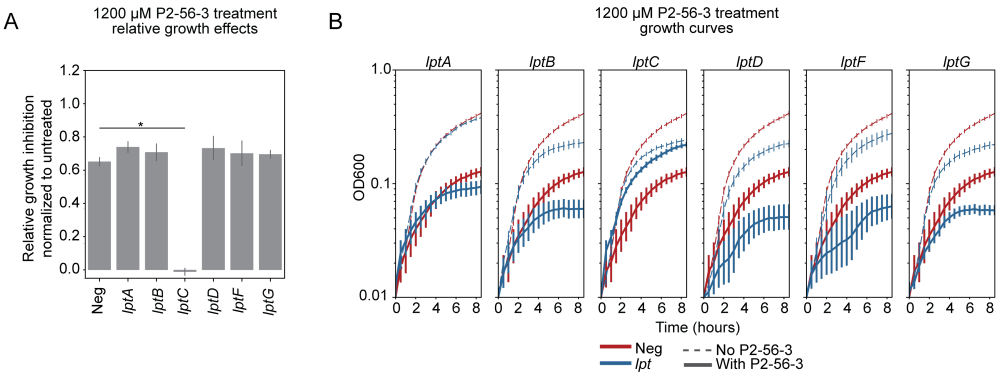

**Fig. S10. Majority of arrayed *lpt* depletion strains show minor, insignificant defects in P2-56-3 only treatment.** (A) Relative growth inhibition at 8 hours and (B) growth curves of LOS transport gene depletions tested with 1200 μM P2-56-3. *: FDR q-value ≤0.05 for comparison between depletion strain and negative (Neg) control. Data shown and error bars represent mean ± SEM (mean of 4 technical replicates and 1 representative biological replicate). P-value and FDR calculations: Materials and Methods.

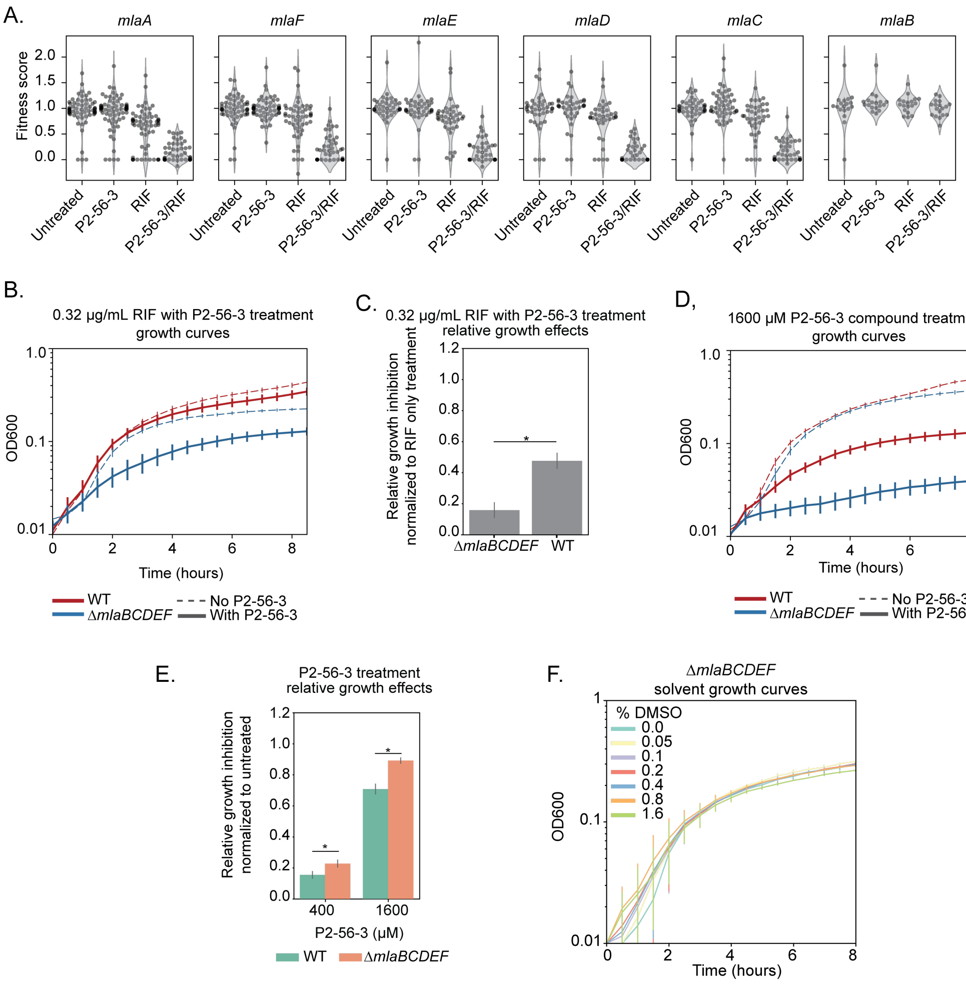

**Fig. S11. MLA complex mutant shows growth inhibitory effects in the presence of P2-56-3.** (A) Fitness effects with all drug treatments across individual MLA complex mutants from nonessential genes screen. (B) Growth curves and (C) relative growth inhibition at 8 hours of MLA knockout mutant and WT treated with 0.32 µg/mL RIF only and in combination with P2-56-3. Data shown and error bars represent mean ± SEM (mean of 4 technical replicates and 2 biological replicates). *: p-value ≤0.05 for comparison between knockout mutant and WT. (D) Growth curves and (E) relative growth inhibition at 8 hours of MLA knockout mutant and WT treated with P2-56-3 only. Data shown and error bars represent mean ± SEM (mean of 4 technical replicates and 3 biological replicates). *: FDR q-value ≤0.05 for comparison between knockout mutant and WT. (F) Growth curves of MLA complex knockout mutant and WT treated with various percentages of

DMSO. Data shown and error bars represent mean ± SEM (mean of 4 technical replicates and 1 representative biological replicate). (C, E) P-value and FDR calculations: Materials and Methods.

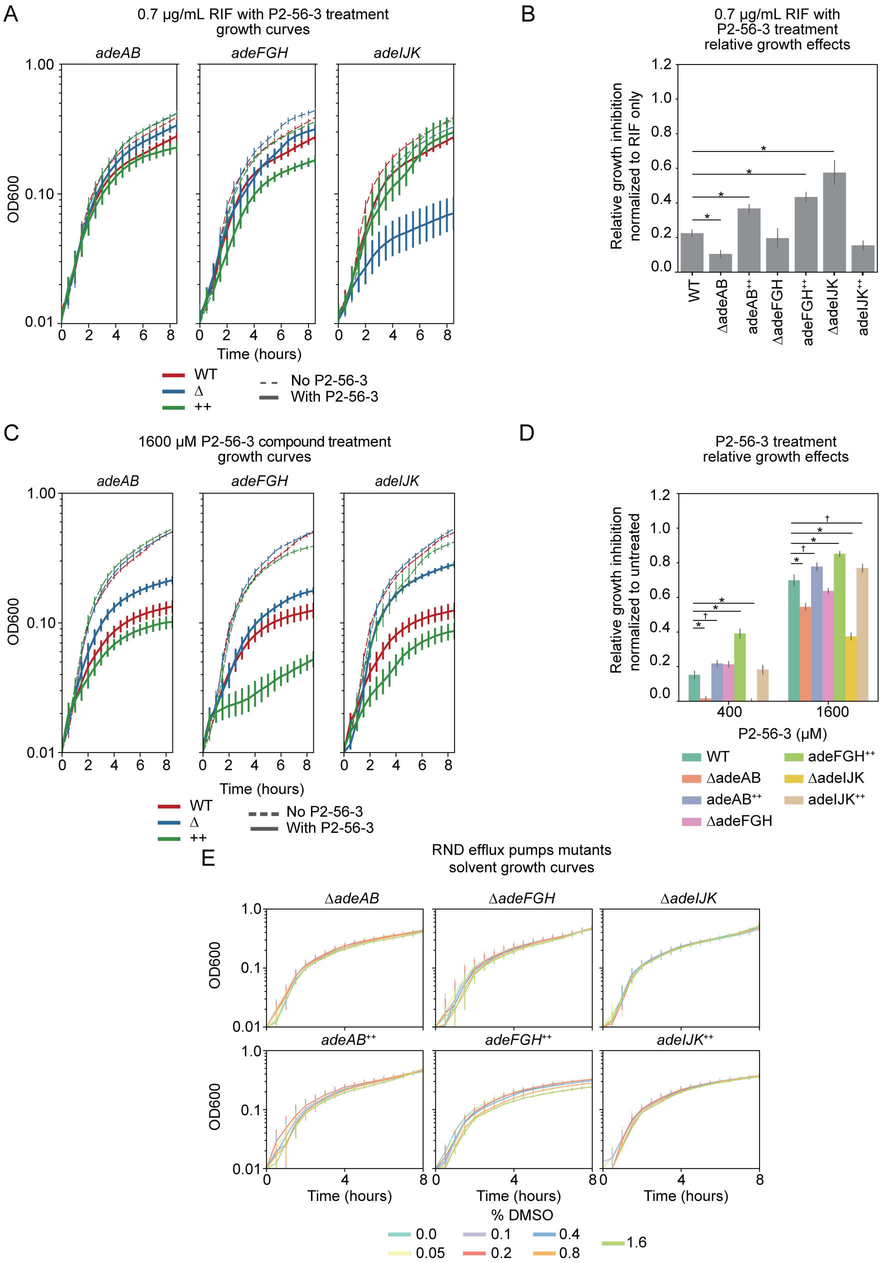

**Fig. S12. RND efflux pump mutants show growth inhibitory effects in the presence of P2-56-3.** (A) Growth curves and (B) relative growth inhibition of RND efflux mutants and WT treated with 0.7 µg/mL RIF only and in combination with P2-56-3. Data shown and error bars represent mean ± SEM (mean of 4 technical replicates and 2 biological replicates). (C) Growth curves and (D) relative growth inhibition at 8 hours of *adeAB, adeFGH,* and *adeIJK* knockout and hyperexpresser strains treated with P2-56-3 only. Data shown and error bars represent mean ± SEM (mean of 4 technical replicates and 2 biological replicates). (E) Growth curves of RND efflux mutants treated with various percentages of DMSO. Data shown and error bars represent the mean ± SEM (mean of 4 technical replicates and 1 representative biological replicate). (B, D) *: FDR q-value ≤0.05 and †: FDR q-value ≤0.1 for comparison between mutant and WT. P-value and FDR calculations: Materials and Methods.

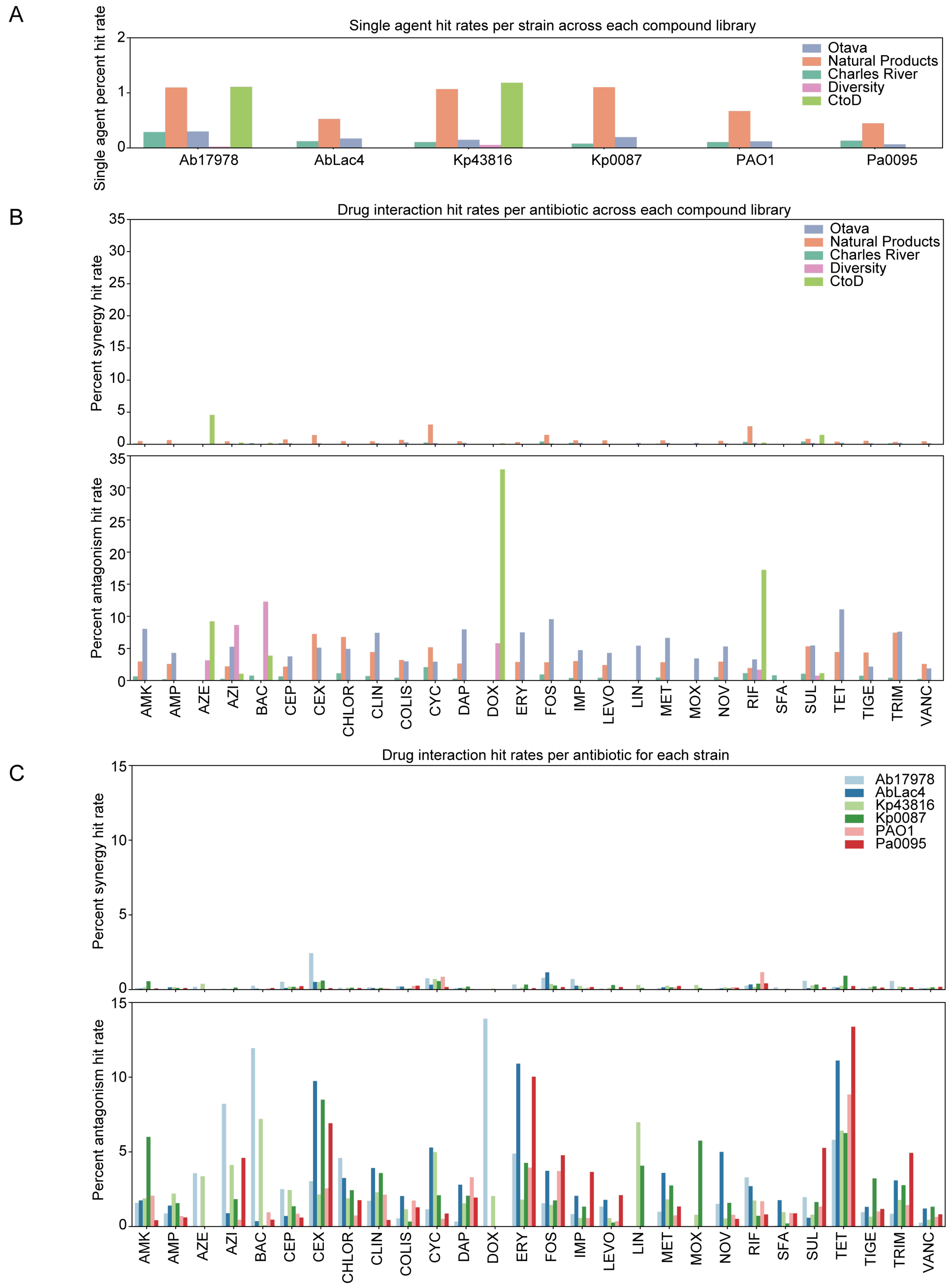

**Fig. S13. Natural products and semi-synthetic compounds have higher hit rates than synthetic compounds.** (A) We calculated single-agent hit rates calculated using the number of compounds that fell above the cutoffs with at least 1 strain (relative growth inhibition ≥0.3 and FDR q-value ≤0.05). We also calculated combination hit rates using the number of strain-antibiotic-compound combinations that fell above or below the cutoffs (top panel, synergy: Bliss sum score

≥0.3 and FDR q-value ≤0.05; bottom panel, antagonism: Bliss sum score ≤-0.3 and FDR q-value ≤0.05) across each (B) compound library and (C) strain. We used the following abbreviations used for each strain: *A. baumannii* ATCC 17978 (Ab17978), *A. baumannii* LAC-4 (AbLac4), *K. pneumoniae* ATCC 43816 (Kp43816), *K. pneumoniae* AR0087 (Kp0087), *P. aeruginosa* PAO1 (PAO1), and *P. aeruginosa* AR0095 (Pa0095). P-value and FDR calculations: Materials and Methods.

**Table S1. Apparent IC90s (µg/mL) of each strain used in the antibiotic combinations screen based on liquid broth cultures in 384W plates.** We determined minimum ICs with growth inhibition values of at least 90%. We estimated IC90s as greater than 2X the maximum tested concentration where we observed inhibition <90% at the maximum tested concentration.

| Antibiotic | Ab17978 | AbLac4 | Kp43816 | Kp0087 | PAO1 | Pa0095 |
|---|---|---|---|---|---|---|
| Amikacin | 0.4 | 9.1 | 2.5 | 0.5 | 4.5 | >10.2 |
| Ampicillin | >256.0 | >512.0 | >512.0 | >512.0 | >512.0 | >512.0 |
| Aztreonam | >128.0 | >324.0 | >0.3 | >512.0 | >8.0 | 51.2 |
| Azithromycin | 1.1 | 9.1 | 64.0 | 51.2 | >16.0 | >10.2 |
| Bactrim | 512.0 | >512.0 | 16.2 | 51.2 | 512 | >512.0 |
| Cefepime | >32.0 | >512.0 | >1.0 | >512.0 | >4.0 | >57.6 |
| Ceftriaxone | >128.0 | >512.0 | >16.0 | >32.0 | >512.0 | >512.0 |
| Chloramphenicol | 51.2 | 287.92 | >512.0 | >512.0 | 512.0 | >57.6 |
| Clindamycin | 91.1 | 287.9 | 512 | 512 | >512.0 | >512.0 |
| Colistin | 0.6 | 1.4 | 3.2 | >512.0 | 3.2 | 2.9 |
| Cycloserine | >512.0 | >512.0 | >512.0 | 512.0 | >512.0 | >512.0 |
| Daptomycin | >512.0 | >512.0 | >512.0 | >512.0 | >512.0 | >512.0 |
| Doxycycline | >32.0 | 22.8 | 5.1 | 91.1 | >64.0 | >18.2 |
| Erythromycin | 7.2 | 91.1 | 512.0 | 512.0 | >128.0 | 91.1 |
| Fosfomycin | >512.0 | 287.9 | >512.0 | >512.0 | >512.0 | 287.9 |
| Imipenem | 2.0 | 3.6 | >16.0 | 8.0 | 1.6 | >18.2 |
| Levofloxacin | 0.2 | 12.8 | >1.0 | 287.9 | .01 | >10.2 |
| Linezolid | 287.9 | >512.0 | >512.0 | >512.0 | >512.0 | >512.0 |
| Metronidazole | >512.0 | >512.0 | >512.0 | >512.0 | >512.0 | >512.0 |
| Moxifloxacin | 0.3 | 7.2 | >2.0 | >512.0 | 1.1 | >18.2 |
| Novobiocin | 10.1 | 40.5 | >512.0 | >512.0 | >512.0 | >512.0 |
| Rifampicin | >32.0 | 5.7 | >128.0 | 45.5 | 36.0 | >32.4 |
| Sulbactam | >256.0 | 51.2 | 256.0 | 81.05 | 512.0 | >512.0 |
| Tetracycline | >64.0 | >512.0 | 8.0 | 512.0 | >32.0 | >18.2 |
| Tigecycline | >128.0 | >256.0 | >64.0 | >512.0 | >512.0 | >182.0 |

| | | | | | | |
|---|---|---|---|---|---|---|
| Trimethoprim | 64.0 | >512.0 | >16.0 | 51.2 | 161.9 | >512.0 |
| Vancomycin | 287.9 | >512.0 | >512.0 | >512.0 | >512.0 | >512.0 |

**Table S2. DropArray antibiotic combinations screening information.** Information about the numbers and statistics associated with the combination screens, discoveryChip dimensions, and antibiotic-antibiotic positive controls. The Charles River library was split up into 2 screening batches as noted below.

| Library | Otava | Natural Products | Charles River (1) | Charles River (2) | Diversity | CtoD |
|---|---|---|---|---|---|---|
| Strains | Ab17978, AbLac4, Kp43816, Kp0087, PAO1, Pa0095 | Ab17978, AbLac4, Kp43816, Kp0087, PAO1, Pa0095 | Ab17978, AbLac4, Kp43816, Kp0087, PAO1, Pa0095 | Ab17978, AbLac4, Kp43816, Kp0087, PAO1, Pa0095 | Ab17978, Kp43816 | Ab17978, Kp43816 |
| Number of antibiotics | 22 | 22 | 21 | 21 | 5 (+1 conc of SUL) | 5 (+1 conc of SUL) |
| Number of concentrations | 3 | 3 | 2 | 2 | 3 | 3 |
| Number of antibiotic concentrations | 66 | 66 | 42 | 42 | 16 | 16 |
| Number of media only inputs | 2 | 3 | 2 | 2 | 4 | 3 |
| Number of bacteria only inputs | 2 | 3 | 2 | 2 | 2 | 2 |
| Number of compounds screened | 1,920 | 800 | 4,734 | 1,030 | 21,089 | 1,236 |
| Number of compounds in combinations (Postfilter) | 1,918 | 791 | 4,726 | 1,030 | 18,439 | 950 |
| Number of chips (post-filter) | 107 | 67 | 297 | 68 | 232 | 28 |
| Number of antibiotic-compound combinations | 189,730 | 91,803 | 557,757 | 122,176 | 199,450 | 9,732 |
| Number of unique antibiotic-compound combinations | 568,393 | 271,427 | 1,115,335 | 243,837 | 531,501 | 25,810 |
| Pre-filter wells | 16,842,168 | 8,717,189 | 44,954,031 | 9,281,668 | 32,761,450 | 2,065,534 |
| Post-filter wells | 11,773,743 | 5,325,573 | 30,573,223 | 5,942,076 | 26,733,450 | 1,456,254 |

| | | | | | | |
|---|---|---|---|---|---|---|
| Percent assays passed filtering | 0.70 | 0.61 | 0.68 | 0.64 | 0.94 | 0.71 |
| Z' passing rate (across all strains) | 0.78 | 0.88 | 0.94 | 1.0 | 0.80 | 0.88 |
| Average antibiotic-compound replicate count | 9 | 10 | 16 | 15 | 18 | 22 |
| Total number of compound inputs per chip | 80 | 64 | 80 | 80 | 80 | 72 |
| Total number of media only inputs per chip | 2 | 3 | 2 | 2 | 4 | 3 |
| Total number of bacteria only inputs per chip | 2 | 3 | 2 | 2 | 4 | 4 |
| Total number of antibiotic inputs per chip | 66 | 66 | 42 | 42 | 32 | 32 |
| Total number of inputs per chip | 150 | 136 | 126 | 126 | 118 | 114 |
| Total number of chips (pre-filter) | 140 | 78 | 336 | 69 | 262 | 32 |
| Average antibiotic-antibiotic replicate count | 12 | 10 | 20 | 22 | 58 | 48 |
| Average antibiotic-media replicate count | 18 | 8 | 13 | 21 | 15 | 61 |
| Average bacteria-compound replicate count | 17 | 9 | 16 | 20 | 17 | 44 |

| **discoveryChip Dimensions** | |
|---|---|
| Number of wells | 139,664 |
| Moat length (µm) | 80 |
| Moat distance above array (µm) | 200 |
| Moat numbers | 50 |
| Well diameter (µM) | 132.6 |

| | | |
|---|---|---|
| Pitch (μm) | 50 | [redacted] |
| Total array footprint (mm) | 73.27 x 110 | [redacted] |
| Well height (μm) | 130-160 | [redacted] |

[redacted]

| Antibiotic-antibiotic synergy controls | | | | |
|---|---|---|---|---|
| **Antibiotic 1** | **Antibiotic 2** | **Screen** | **Strain** | [redacted] |
| SUL | IMP | Charles River (1) | Ab17978 | [redacted] |
| SUL | IMP | Charles River (1) | AbLac4 | [redacted] |
| TRIM | SFA | Charles River (1) | Kp43816 | [redacted] |
| AMP | SUL | Charles River (1) | Kp0087 | [redacted] |
| SUL | IMP | Charles River (1) | PAO1 | [redacted] |
| TRIM | SFA | Charles River (1) | Pa0095 | [redacted] |
| SUL | IMP | Charles River (2) | Ab17978 | [redacted] |
| SUL | IMP | Charles River (2) | AbLac4 | [redacted] |
| TRIM | SFA | Charles River (2) | Kp43816 | [redacted] |
| AMP | SUL | Charles River (2) | Kp0087 | [redacted] |
| TRIM | SFA | Charles River (2) | PAO1 | [redacted] |
| TRIM | SFA | Charles River (2) | Pa0095 | [redacted] |
| CEX | SUL | Natural Products | Ab17978 | [redacted] |
| CEX | SUL | Natural Products | AbLac4 | [redacted] |
| AMP | SUL | Natural Products | Kp43816 | [redacted] |
| LEVO | RIF | Natural Products | Kp0087 | [redacted] |
| AMP | SUL | Natural Products | PAO1 | [redacted] |
| SUL | IMP | Natural Products | Pa0095 | [redacted] |

| | | | | | |
|---|---|---|---|---|---|
| RIF | COLIS | Otava | Ab17978 | [redacted] | [redacted] |
| FOS | TRIM | Otava | AbLac4 | [redacted] | [redacted] |
| SUL | AMP | Otava | Kp43816 | [redacted] | [redacted] |
| SUL | CEP | Otava | Kp0087 | [redacted] | [redacted] |
| SUL | IMP | Otava | PAO1 | [redacted] | [redacted] |
| SUL | IMP | Otava | Pa0095 | [redacted] | [redacted] |
| AZE | SUL | Diversity | Ab17978 | [redacted] | [redacted] |
| AZE | SUL | Diversity | Kp43816 | [redacted] | [redacted] |
| AZE | SUL | CtoD | Ab17978 | [redacted] | [redacted] |
| AZE | SUL | CtoD | Kp43816 | [redacted] | [redacted] |

**Table S3. Selected compound potentiator hits and their associated hit profile.** For each compound, the compound library and strain-antibiotic interaction the compound scored with was noted for the primary screen and DropArray checkerboards. For checkerboards, Bliss sum scores were calculated. If combinations exhibited checkerboard Bliss sum scores ≥0.3, we calculated FIC scores and noted those with $FIC_{min}$ ≤0.5 as a supplemental measure of their synergistic effects.

| Compound name | Library | Strain-antibiotic primary screen hit | Strain-antibiotic checkerboard Bliss sum hit (Bliss sum ≥0.3) | Strain-antibiotic checkerboard FIC hit ($FIC_{min}$ ≤0.5) |
|---|---|---|---|---|
| BRD0433 | Diversity | Kp43816-RIF | Kp43816-RIF, Kp0087-RIF | Kp43816-RIF |
| IIIA:8:G | CtoD | Ab17978-RIF, Kp43816-AZI, Kp43816-RIF, Kp43816-SUL | Ab17978-RIF, Ab17978-ERY, Kp43816-AZE, Kp43816-AZI, Kp43816-DOX, Kp43816-ERY, Kp43816-RIF, Kp43816-TIGE, Kp0087-AZE, Kp0087-AZI, Kp0087-BAC, Kp0087-DOX, Kp0087-ERY, Kp0087-RIF, Kp0087-TIGE, Kp0087-VANC | Kp43816-AZI, Kp43816-BAC, Kp4386-DOX, Kp43816-ERY, Kp43816-RIF, Kp43816-SUL, Kp0087-AZI, Kp0087-RIF, Kp0087-TIGE |
| BRD1479 | Otava | Kp43816-TRIM, PAO1-TRIM | Kp0087-NOV, Kp0087-SUL, PAO1-CYC, PAO1-TRIM, Pa0095-TRIM | PAO1-CYC[a], PAO1-TRIM[a], Pa0095-TRIM[a] |
| BRD4550 | Otava | Ab17978-CEP | Ab17978-CEP, Kp43816-FOS, PAO1-AMP, PAO1-CEP, PAO1-FOS, PAO1-IMP, PAO1-SUL, Pa0095-TIGE, Pa0095-TRIM | Ab17978-CEP, PAO1-IMP |
| Esculin Monohydrate | Natural Products | Kp0087-AZI, Kp0087-CLIN, Kp0087-ERY, Kp0087-CHLOR, Kp0087-RIF | Kp0087-AZI, Kp0087-CHLOR, Kp0087-CLIN, Kp0087-COLIS, Kp0087-ERY | Kp0087-AZI, Kp0087-ERY |
| P2-56 | Otava | Ab17978-COLIS, Ab17978-NOV, Ab17978-RIF, Ab17978-SUL, AbLac4-CLIN, AbLac4-NOV, AbLac4-RIF, Kp43816-CLIN, Kp43816-LIN, Kp43816-RIF, Kp0087-NOV, Pa0095-TET | Ab17978-NOV, Ab17978-RIF, Kp0087-LEVO, Kp0087-MOX, PAO1-CYC, Pa0095-TET | AbLac4-NOV, Kp43816-RIF, Kp0087-MOX, Pa0095-TET |

[a]FIC50s were calculated instead of FIC90s

**Table S4. All strains and plasmids used in this study.** Strains used in the screens, structure-activity-relationship studies, checkerboard assays, and knockout and knockdown generation with associated plasmids.

| Strain or plasmid | Genotype or description | Reference |
|---|---|---|
| | *E. coli* | |
| DH5α | supE44 DlacU169 (f80lacZDM15) hsdR17 recA1 endA1 gyrA96 thi-1 relA1 | (11) |
| DH5α λpir | DH5α λpir tet::Mu recA | (12) |
| K-12 | K-12 substr. MG1655 | (13) |
| | *K. pneumoniae* | |
| ATCC 43816 | ATCC 43816 | |
| AR 0087 | Unknown, Biosample Accession #: SAMN04014928, CDC AR Isolate Bank | (14) |
| | *P. aeruginosa* | |
| PAO1 | wound isolate | (15) |
| AR 0095 | Unknown, Biosample Accession #: SAMN04014936, CDC AR Isolate Bank | (14) |
| | *S. aureus* | |
| Newman | human infection isolate | (16) |
| | *E. faecium* | |
| ATCC BAA-2317 | ATCC BAA-2317, human feces | |
| | *A. baumannii* | |
| ATCC 17978 | ATCC 17978, cerebrospinal fluid isolate | |
| AB5075 | bone isolate/osteomyelitis | (17) |
| EGA355 | sputum isolate | Provided by Ralph Isberg's lab |
| EGA366 | urine isolate | Provided by Ralph Isberg's lab |
| EGA368 | sputum isolate | Provided by Ralph Isberg's lab |
| LAC-4 | Outbreak strain isolated from hospital A in 1997 | (18) |

| | | |
|---|---|---|
| *ΔmlaBCDEF* | ATCC 17978 ΔmlaBCDEF | Provided by Ralph Isberg's lab |
| *ΔadeAB* | ATCC 17978 ΔadeAB | This study |
| *ΔadeFGH* | ATCC 17978 adeLΔI335A336, ΔadeFGH | Provided by Ralph Isberg's lab |
| *ΔadeIJK* | ATCC 17978 ΔadeIJK | Provided by Ralph Isberg's lab |
| *adeAB*++ | ATCC 17978 adeS (R152S) | Provided by Ralph Isberg's lab |
| *adeFGH*++ | ATCC 17978 adeLΔI335A336 | Provided by Ralph Isberg's lab |
| *adeIJK*++ | ATCC 17978 adeN :: Isaba1 | Provided by Ralph Isberg's lab |
| EHA136 | ATCC 17978 attTn7::tetR-tetP-dcas9-rrnBT1-T7Te Gmr | Provided by Ralph Isberg's lab |
| *lptA* | EHA136 with sgRNA (ACX60_RS11965_17) | This study |
| *lptB* | EHA136 with sgRNA (ACX60_RS11960_1) | This study |
| *lptC* | EHA136 with sgRNA (ACX60_RS11970_23) | This study |
| *lptD* | EHA136 with sgRNA (ACX60_RS10360_20) | This study |
| *lptF* | EHA136 with sgRNA (ACX60_RS16930_6) | This study |
| *lptG* | EHA136 with sgRNA (ACX60_RS16925_4) | This study |
| Negative | EHA136 with sgRNA (non-targeting control) | Provided by Ralph Isberg's lab |
| | Plasmids | |
| pWH1266 | ori pBR322 ori pWH1277 MCS Tcr Cbr | (19) |
| pJB4648 | GmR derivative of pSR47S | (20) |
| pBBR1(MCS5)-Plac-gfp | Constitutive GFP GmR | Provided by Katharina Ribbeck's lab |
| pACYC184-GFP | Consitutive GFP CmR | Provided by Joan Mecsas's lab |
| pEGE305-GFP | IPTG-inducible GFP TetR | Provided by Ralph Isberg's lab |
| pEGE9 - GFP | Consitutive GFP CarbR | Provided by Ralph Isberg's lab |

**Table S5. All oligonucleotides used in this study.** Oligonucleotides used in the generation of CRISPRi and Tn-seq screens, as well as knockout and knockdown mutant generation.

| **Cloning** |
|---|

| Primer name | Sequence (5' - 3'; restriction site underlined) | RE site(s) | Description | Gene |
|---|---|---|---|---|
| Gene deletion | | | | |
| MT_o021 | acccggGTCGACcttacttacatggctatctac | SalI | adeAB Upstream 750bp Forward | ACX60_RS09125, ACX60_RS09130 |
| MT_o022 | ctgcagGGTACCgtccaaacctagtgagtttttg | KpnI | adeAB Upstream 6nt Reverse | ACX60_RS09125, ACX60_RS09130 |
| MT_o025 | ctgcagGGTACCggaaatactcaatttcctctgatgc | KpnI | adeAB Downstream 6nt Forward | ACX60_RS09125, ACX60_RS09130 |
| MT_o026 | acccggGGATCCcaaataaagtaagtgccaagttc | BamHI | adeAB Downstream 750bp Reverse | ACX60_RS09125, ACX60_RS09130 |
| Gene knockdown | | | | |
| MT_d077 | TAGTGAAGAACCTTTGAAATTAAGACTT | BsaI | lptA_17 | ACX60_RS11965 |
| MT_d087 | TAGTTTGCTGCATTAGTCACGCGATCCT | BsaI | lptB_1 | ACX60_RS11960 |
| MT_d091 | TAGTTATAAAACTCAGGTATCCTTGAGC | BsaI | lptC_23 | ACX60_RS11970 |
| MT_d105 | TAGTGAGGGTAAAAATGGCGGTCGCTAA | BsaI | lptD_20 | ACX60_RS10360 |
| MT_d109 | TAGTAATAATCAAATTAATATTTCCAAA | BsaI | lptF_6 | ACX60_RS16930 |
| MT_d119 | TAGTGCGCAGTGGTTTTCGTCACAAATT | BsaI | lptG_4 | ACX60_RS16925 |
| MT_d078 | AAACAAGTCTTAATTTCAAAGGTTCTTC | BsaI | lptA_17 | ACX60_RS11965 |
| MT_d088 | AAACAGGATCGCGTGACTAATGCAGCAA | BsaI | lptB_1 | ACX60_RS11960 |
| MT_d092 | AAACGCTCAAGGATACCTGAGTTTTATA | BsaI | lptC_23 | ACX60_RS11970 |

| | | | | |
|---|---|---|---|---|
| MT_d106 | AAACTTAGCGACCGCCATTTTTACCCTC | BsaI | lptD_20 | ACX60_RS10360 |
| MT_d110 | AAACTTTGGAAATATTAATTTGATTATT | BsaI | lptF_6 | ACX60_RS16930 |
| MT_d120 | AAACAATTTGTGACGAAAACCACTGCGC | BsaI | lptG_4 | ACX60_RS16925 |
| Neg7_F | TAGTTGATCCAATACTATGTAACACTAA | BsaI | Non-targeting guide | |
| Neg7_R | AAACTTAGTGTTACATAGTATTGGATCA | BsaI | Non-targeting guide | |

| **Mariner Tn-seq sequencing library construction** | | |
|---|---|---|
| Primer Name | Sequence (5' - 3') | Index |
| First PCR | | |
| Nextera 2A-R | GTCTCGTGGGCTCGGAGATGTGTATAAGAGACAG | |
| olj638 | CTGTGTGGGCACTCGATGACGTCAGACC | |
| Second PCR -- Rightward Nextera Indexed primers | | |
| olk141 | CAAGCAGAAGACGGCATACGAGATCCGCCTGCGTCTCGTGGGCTCGGAGATGTG | GCAGGCGG |
| N703 | CAAGCAGAAGACGGCATACGAGATTTCTGCCTGTCTCGTGGGCTCGGAGATGTG | AGGCAGAA |
| N708 | CAAGCAGAAGACGGCATACGAGATCCTCTCTGGTCTCGTGGGCTCGGAGATGTG | CAGAGAGG |
| N710 | CAAGCAGAAGACGGCATACGAGATCAGCCTCGGTCTCGTGGGCTCGGAGATGTG | CGAGGCTG |
| N711 | CAAGCAGAAGACGGCATACGAGATTGCCTCTTGTCTCGTGGGCTCGGAGATGTG | AAGAGGCA |
| Second PCR - Leftward mariner specific Indexed primers | | |
| mar147 | AATGATACGGCGACCACCGAGATCTACACGCAGGCGGCGTTGACCGGGGACTTATCAGCCAACCTGTTA | GCAGGCGG |
| mar148 | AATGATACGGCGACCACCGAGATCTACACAGGCAGAACGTTGACCGGGGACTTATCAGCCAACCTGTTA | AGGCAGAA |

| | | |
|---|---|---|
| mar149 | AATGATACGGCGACCACCGAGATCTACACCAGAGAG GCGTTGACCGGGGACTTATCAGCCAACCTGTTA | CAGA GAGG |
| mar150 | AATGATACGGCGACCACCGAGATCTACACCGAGGCT GCGTTGACCGGGGACTTATCAGCCAACCTGTTA | CGAG GCTG |
| mar151 | AATGATACGGCGACCACCGAGATCTACACAAGAGGCA CGTTGACCGGGGACTTATCAGCCAACCTGTTA | AAGA GGCA |
| mar152 | AATGATACGGCGACCACCGAGATCTACACGAGGAGC CCGTTGACCGGGGACTTATCAGCCAACCTGTTA | GAGG AGCC |
| | Reconditioning | |
| P1 | AATGATACGGCGACCACCGA | |
| P2 | CAAGCAGAAGACGGCATACGA | |
| | Sequencing | |
| mar512 | CGTTGACCGGGGACTTATCAGCCAACCTGTTA | |

| **CRISPRi sequencing library construction** | |
|---|---|
| Primer Name | Sequence (5' - 3') |
| | PCR1 |
| Primer For2_sgRNA_Pool | TCGTCGGCAGCGTCAGATGTGTATAAGAGACAG GTCCTGTGGATCCGATCTTTGAC |
| Primer Rev_sgRNA_Pool | GTCTCGTGGGCTCGGAGATGTGTATAAGAGACAG AGATGAGTTTTTGTTCGGGCCC |
| | PCR2 |
| Nextera N5x index Primer | AATGATACGGCGACCACCGAGATCTACAC [i5] TCGTCGGCAGCGTC |
| Nextera N7x index Primer | CAAGCAGAAGACGGCATACGAGAT [i7] GTCTCGTGGGCTCGG |
| | i5 Nextera Primers |
| N503 | TATCCTCT |
| N505 | GTAAGGAG |
| N506 | ACTGCATA |

| | | |
|---|---|---|
| N507 | AAGGAGTA | [redacted] |
| i7 Nextera Primers | | |
| N701 | TAAGGCGA | |
| N702 | CGTACTAG | |
| N703 | AGGCAGAA | |
| N704 | TCCTGAGC | |
| N705 | GGACTCCT | |
| N706 | TAGGCATG | |
| N708 | CAGAGAGG | |
| N709 | GCTACGCT | |
| N711 | AAGAGGCA | |

**Table S6. All antibiotics and concentrations used in the primary screens.** Antibiotic shorthand is the abbreviation used for the antibiotics in the dataset, linked to the antibiotic name, manufacturer and catalog number. Additionally, we show the stock concentrations made (mg/mL) and the solvent the powders were dissolved in.

| Antibiotic shorthand | Antibiotic name | Company | Catalog number | Stock concentration (mg/mL) | Solvent |
|---|---|---|---|---|---|
| AMP | Ampicillin | AmericanBio | AB00115 | 50 | H2O with 0.1% Triton X-100 |
| AMK | Amikacin | Toronto Research Chemicals | A578490 | 15 | H2O with 0.1% Triton X-100 |
| AMX | Amoxicillin | Aurum | GN63361 | 25 | DMSO |
| AZE | Aztreonam | AstaTech | 42414 | 30 | DMSO |
| CEX | Cefoxitin | Combi-Blocks | QC-6999 | 50 | DMSO |
| CHLOR | Chloramphenicol | Sigma-Aldrich | C0376 | 50 | DMSO |
| CLIN | Clindamycin | Sigma-Aldrich | PH011629 | 50 | DMSO |
| COLIS | Colistin | Sigma-Aldrich | C4461 | 25 | H2O with 0.1% Triton X-100 |
| CYC | Cycloserine | Sigma-Aldrich | C6880 | 20 | H2O with 0.1% Triton X-100 |
| DAP | Daptomycin | Combi-Blocks | QC-4306 | 50 | DMSO |
| DOX | Doxycycline | eNovation | K35178 | 10 | H2O with 0.1% Triton X-100 |
| ERY | Erythromycin | Sigma-Aldrich | E5589 | 20 | DMSO |
| FOS | Fosfomycin | MP Biomedicals | 151876 | 25 | H2O with 0.1% Triton X-100 |
| LEVO | Levofloxacin | AstaTech | 31131 | 5 | DMSO |
| LIN | Linezolid | AstaTech | 25787 | 25 | DMSO |
| MEC | Mecillinam | Cayman Chemical Company | 9002008 | 50 | H2O with 0.1% Triton X-100 |
| MET | Metronidazole | AstaTech | 28634 | 50 | DMSO |

| | | | | | |
|---|---|---|---|---|---|
| MOX | Moxifloxacin | Combi-Blocks | QB-4316 | 15 | DMSO |
| NOV | Novobiocin | Sigma-Aldrich | 74675 | 20 | H2O with 0.1% Triton X-100 |
| RIF | Rifampicin | TCI Chemicals | R0079 | 15 | DMSO |
| SUL | Sulbactam | AstaTech | 37516 | 25 | H2O with 0.1% Triton X-100 |
| SFA | Sulfamethoxazole | AmBeed | A205081 | 50 | DMSO |
| TET | Tetracycline | Sigma-Aldrich | T7660 | 50 | DMSO |
| TIGE | Tigecycline | AstaTech | 25103 | 50 | DMSO |
| TRIM | Trimethoprim | AstaTech | 28264 | 50 | DMSO |
| VANC | Vancomycin | Sigma-Aldrich | V2002 | 50 | H2O with 0.1% Triton X-100 |
| AZI | Azithromycin | AstaTech | N46072 | 15 | DMSO |
| CEP | Cefepime | eNovation Chem | D371159 | 25 | DMSO |
| IMP | Imipenem | Aurum | X-2302 | 5 | H2O with 0.1% Triton X-100 |
| BAC | Bactrim (made by mixing 1:5 ratio of TRIM and SFA) | -- | -- | 50 | DMSO |

**SI Datasets. Available for access on the Harvard Dataverse network (https://dataverse.harvard.edu/privateurl.xhtml?token=23f475d0-c94e-4ad7-aa2b-2cac89c6196e).**

**Dataset S1 (separate file). All individual Bliss scores for each antibiotic compound interaction at different antibiotic concentrations from the DropArray screen.** Concentrations used per screen in µg/mL. These are linked to the labels used in the dataset, where a corresponds to the highest concentration. Sheet 1 - screen_name: name of the compound library, strain_name: name of the bacterial strain used, abx_name: antibiotic shorthand name, abx_conc: antibiotic concentration used in µg/mL, cp_name: name of the compound, bliss_med: Bliss score calculated for the combination, bliss_se: Bliss standard error calculated via bootstrapping, Ea_med: antibiotic growth inhibitory effect, Ea_se: standard error of the antibiotic growth inhibitory effect, Ec_med: compound growth inhibitory effect, Ec_se: standard error of the compound growth inhibitory effect, Eac_med: antibiotic and compound growth inhibitory effect together, Eac_se: standard error of the antibiotic and compound growth inhibitory effects, cp_SMILES: SMILES string associated with the compound. Sheet2 - screen: compound library, abx_strain_letconc: antibiotic name, strain name, and letter concentration used where a is the highest and c is the lowest concentration, conc: concentration in µg/mL.

**Dataset S2 (separate file). All Bliss sum scores for each antibiotic compound interaction from the DropArray screen.** screen_name: name of the compound library which is associated with different batches of screening, strain_name: name of the bacterial strain used, abx_name: antibiotic shorthand name, cp_name: name of the compound, bliss_med: Bliss score summed across the antibiotic concentrations tested in combination with compound, bliss_se: sum of the Bliss standard errors across the antibiotic concentrations tested in combination with compound, cp_SMILES: SMILES string associated with the compound, c_batch_bliss_pval: p-values calculated on a per screen basis, c_adj_batch_bliss_pval: FDR q-value calculated on a per screen basis.

**Dataset S3 (separate file). All compound only growth inhibition data from the DropArray screen per strain.** cp_name: name of the compound, norm_growth_inh: growth inhibitory effect of the compound, norm_growth_SE: standard error of the growth inhibitory effect of the compound, adj_batchpval: FDR q-value calculated per screen, screen_name: name of the compound library which is associated with different batches of screening, strain_name: name of the bacterial strain used, cp_SMILES: SMILES string associated with the compound.

**Dataset S4 (separate file). Individual and aggregated fitness scores across all guides in the essential genes screen across all treatment conditions.** Sheet 1 - Guide Name: gene targeted with associated guide number label, Gene Target: gene name, Description: description of the protein product, First Nucleotide # Mutated Relative to PAM: position of the nucleotide in the guide that is first mutated relative to the location of the PAM site, Second Nucleotide # Mutated Relative to PAM: position of the nucleotide in the guide that is mutated additionally relative to the location of the PAM site, ATCC_Locus: locus tags using NDEONHPJ labels, ACX_Locus: locus tags using ACX60_RS labels, d_R0C100_t2: fitness difference of the compound treated relative to the untreated, induced populations, p_R0C100_t2: p-value of the compound treated relative to the untreated, induced populations, q_R0C100_t2: FDR q-value of the compound treated relative to the untreated, induced populations, d_R7C0_t2: fitness difference of the rifampin treated relative to the untreated, induced populations, p_R7C0_t2: p-value of the rifampin treated relative to the untreated, induced populations, q_R7C0_t2: FDR q-value of the rifampin treated relative to the untreated, induced populations, d_R7C100_t2: fitness difference of the rifampin and compound treated relative to the untreated, induced populations, p_R7C100_t2: p-value of the rifampin and compound treated relative to the untreated, induced populations, q_R7C100_t2: FDR q-value of the rifampin and compound treated relative to the untreated, induced populations, d_R0C0_t1: fitness difference of the compound population relative to the untreated, induced population, p_R0C0_t1: p-value of the compound population relative to the untreated, induced population, q_R0C0_t1: FDR q-value of the compound population relative to the untreated, induced population, d_R7C100vsR7C0: fitness difference of the rifampin and compound treated relative to the rifampin, induced population, p_R7C100vsR7C0: p-value of the rifampin and compound treated relative to the rifampin, induced populations, q_R7C100vsR7C0: FDR q-value of the rifampin and compound treated relative to the rifampin, induced populations, neg: labeled as negative, non-targeting controls. Sheet 2: The same labels apply here except the p-value and FDR q-value were calculated over the aggregated guides. The fitness differences were calculated over the means of the aggregated guides.

**Dataset S5 (separate file). Aggregated fitness scores across the transposon mutants in the nonessential genes screen across all treatment conditions.** ATCC_Locus: locus tags using NDEONHPJ labels, ACX_Locus: locus tags using ACX60_RS labels, Untreated_W: average fitness score of the untreated, Untreated_Count: number of mutants in the untreated group, Untreated_SD: standard deviation of the fitness scores of the untreated, Untreated_SE: standard error of the fitness scores of the untreated, R0C100_W: average fitness score of the lower compound only treatment, R0C100_Count: number of mutants in the lower compound only group, R0C100_SD: standard deviation of the lower compound only treatment, R0C100_SE: standard error of the lower compound only treatment, R7C0_Count: number of mutants in the rifampin only group, R7C0_SD: standard deviation of the rifampin only treatment, R7C0_SE: standard error of the rifampin treatment, R7C100_W: fitness of the rifampin and compound treatment, R7C100_Count: number of mutants in the rifampin and compound group, R7C100_SD: standard deviation of the rifampin and compound treatment, R7C100_SE: standard error of the rifampin and compound treatment, p_R0C100: p-value of the lower compound only treatment relative to untreated, d_R0C100: fitness difference of lower compound only treatment relative to untreated, q_R0C100: FDR q-value of the lower compound only treatment relative to untreated, p_R7C0: p-value of the rifampin only treatment relative to untreated, d_R7C0: fitness difference of rifampin only treatment relative to untreated, q_R7C0: FDR q-value of the rifampin only treatment relative to untreated, p_R7C100: p-value of the rifampin and compound treatment relative to untreated, d_R7C100: fitness difference of rifampin and compound treatment relative to untreated, q_R7C100: FDR q-value of the rifampin and compound treatment relative to untreated, p_R7C100vsR7C0: p-value of the lower compound only treatment relative to untreated, d_R7C100vsR7C0: fitness difference between the rifampin and compound treatment relative to untreated, q_R7C100vsR7C0: FDR q-value of the rifampin and compound treatment relative to rifampin, Contig: location of gene, Gene: gene name if available, Product: description of the protein product.

**Dataset S6 (separate file). Single agent hits and chemical properties evaluation from the DropArray screen.** Sheet1 contains the growth inhibitory compound hit rates on a screen and strain basis. Sheet2 contains the properties associated with the relative growth inhibition hits compared to the rest of the non-hit compounds. Sheet 1 - hit-num: number of hits that met the criteria (FDR q-value ≤0.05 and relative growth inhibition ≥0.3), percent-hit-rate: number of hits that met the criteria divided by the total number of strain-compounds tested, total-num-combinations: total number of strain-compounds screened for each library and strain. Sheet 2 - adj-p: FDR q-value of hit set vs. the non-hit set property values, diff-mean: mean of the hit set minus the mean of the non-hit set property values, hit-std: standard deviation of the hit set property values, lib-std: standard deviation of the non-hit set property values, hit-mean: mean of the hit set property values, hit-lib: mean of the non-hit set property values, hit-max: max of the hit set property values, hit-min: min of the hit set property values, lib-max: max of the non-hit set property values, lib-min: min of the non-hit set property values, lib-count: number of compounds in the non-hit set, hit-count: number of compounds in the hit set.

**Dataset S7 (separate file). Drug interaction rates and chemical properties evaluation from the DropArray screen.** Sheet1 contains the combination hit rates for each antibiotic-compound-strain combination on a per strain, antibiotic, and screen basis. Sheet2 contains the properties associated with the synergy and antagonism hits compared to the rest of the non-hit compounds. Sheet 1 - syn_hit-num: number of synergistic hits that met the criteria (FDR q-value ≤0.05 and Bliss sum score ≥0.3), syn_percent-hit-rate: number of synergistic hits that met the criteria divided by the total number of combinations, ant_hit-num: number of antagonistic hits that met the criteria

(FDR q-value ≤0.05 and Bliss sum score ≤-0.3) , ant_percent-hit-rate: number of antagonistic hits that met the criteria divided by the total number of combinations, total-num-combinations: total number of strain-antibiotic-compound combinations screened for each library, strain, and antibiotic set. Sheet 2 - ant_adj-p: FDR q-value of antagonistic hit set vs. the non-hit set property values, ant_diff-mean: mean of the antagonistic hit set minus the mean of the non-hit set property values, ant_hit-std: standard deviation of the antagonistic hit set property values, ant_lib-std: standard deviation of the non-antagonistic set property values, ant_hit-mean: mean of the antagonistic hit set property values, ant_hit-lib: mean of the non-antagonistic set property values, ant_hit-max: max of the antagonistic hit set property values, ant_hit-min: min of the antagonistic hit set property values, ant_lib-max: max of the non-antagonistic set property values, ant_lib-min: min of the non-antagonistic set property values, ant_lib-count: number of compounds of the non-antagonistic set, ant_hit-count: number of compounds of the antagonistic hit set, syn_adj-p: FDR q-value of synergistic hit set vs. the non-hit set property values, syn_diff-mean: mean of the synergistic hit set minus the mean of the non-hit set property values, syn_hit-std: standard deviation of the synergistic hit set property values, syn_lib-std: standard deviation of the non-synergistic set property values, syn_hit-mean: mean of the synergistic hit set property values, syn_hit-lib: mean of the non-synergistic set property values, syn_hit-max: max of the synergistic hit set property values, syn_hit-min: min of the synergistic hit set property values, syn_lib-max: max of the non-synergistic set property values, syn_lib-min: min of the non-synergistic set property values, syn_lib-count: number of compounds of the non-synergistic set, syn_hit-count: number of compounds of the synergistic hit set.
